# Remembered event features shape default mode network engagement during emotional memory recall

**DOI:** 10.64898/2026.05.05.721961

**Authors:** Nina Curko, Rosalie Samide, Valentina Krenz, Elizabeth A. Kensinger, Maureen Ritchey

## Abstract

Episodic memory involves reconstruction of past events through integration of multiple types of information, including perceptual details, narrative content, and emotional tone. The default mode network (DMN) is a set of regions thought to support episodic retrieval, yet it remains unclear how distinct subnetworks contribute to recall of these different memory features. Here, we examined how DMN subnetworks support and represent memory for naturalistic emotional events. Male and female human participants encoded short news videos that varied in emotional valence and recalled them in response to neutral cues during functional MRI scanning. Videos were again recalled one day later and memories were scored for perceptual and narrative details. Activity in the dorsomedial subnetwork was related to the emotional valence of the memory, while activity in the medial temporal subnetwork was associated with the number of perceptual details recalled. Multivariate pattern analyses further revealed that the medial temporal subnetwork exhibited greater pattern stability across recalls when recalling more perceptual details, while stability in the dorsomedial and core subnetworks was tied to emotional remembering. Our findings suggest that the dorsomedial subnetwork provides an affective frame for a memory, while the medial temporal subnetwork contributes perceptual specificity. These results demonstrate that the contents of memory retrieval shape network engagement during emotional recall, providing insight into how the brain reconstructs complex real-world experiences.

**Significance Statement:** How do we remember the emotional tone of an event versus its visual details? This study examines how distinct subnetworks within the brain’s default mode network (DMN) contribute to remembering different features of memory. While the medial temporal subnetwork is connected to retrieving the perceptual details of past events, the dorsomedial subnetwork supports recall of the emotional tone. Furthermore, patterns of activity in the medial temporal subnetwork are more stable when recalling more perceptually rich memories, while stability in the dorsomedial subnetwork is tied to emotional remembering. These findings suggest that the default mode network flexibly responds to different kinds of memory features, supporting the reconstruction of rich emotional memories from complex real-world experiences.

## Introduction

Episodic memory retrieval involves reconstructing multiple event features into a cohesive representation. This process is thought to be supported by the default mode network (DMN), a set of interconnected regions active during internal cognitive processes such as memory retrieval, episodic construction, and emotional experiences (Andrews-Hanna et al., 2010; Buckner et al., 2008; Rugg & Vilberg, 2013; Satpute & Lindquist, 2019). Emerging evidence suggests the DMN represents and integrates event-specific information in memory (Ritchey & Cooper, 2020). A major factor affecting how event features contribute to the experience of memory is the presence of positive or negative emotional content (Williams et al., 2022). However, it remains unclear whether different DMN components contribute to remembering distinct event features, and in particular, how DMN engagement during recall is shaped by the emotionality of memory.

The DMN comprises functionally distinct subnetworks, including a medial temporal subsystem associated with episodic processes and a dorsomedial subsystem associated with social cognition (Andrews-Hanna et al., 2010). Within the context of memory, these subnetworks may be differently related to the content of event representations, with regions in the medial temporal subsystem supporting memory for spatial and event context (Ranganath & Ritchey, 2012) and regions in the dorsomedial subsystem supporting representations of objects and their social and emotional associations (Barnett et al., 2021; Karagoz et al., 2023). These networks are differentially recruited when people orient toward the perceptual details and conceptual themes of memory (Ferris et al., 2025; Gurguryan & Sheldon, 2019). These frameworks suggest flexible recruitment of these subnetworks depending on memory content, and specifically, the perceptual and narrative richness of recall.

Another open question is how DMN subnetworks contribute to emotional memories. Emotional content biases memory toward central event details (Kensinger et al., 2007), facilitating abstraction of an event’s high-level meaning. This abstraction process has been attributed to the dorsomedial prefrontal cortex (Baetens et al., 2014), implicating this region – and its associated DMN subnetwork – in integrating an event’s affective tone (Kensinger & Ford, 2021). For real-world memories, both perceptual and narrative details may contribute to affective experience; however, they may be differently weighted for negative and positive valenced memories, with stronger engagement of perceptual regions during negative memory retrieval (Erk et al., 2003; Markowitsch et al., 2003; Mickley & Kensinger, 2008). Together, these findings suggest that emotion modulates how DMN subnetworks contribute to the reconstruction of different event details, but this has not been explicitly tested.

To address these questions, this study aims to characterize the role of DMN subnetworks in retrieving the details of emotional and neutral memories. Video stimuli were used to promote rich, varied recall of life-like events. Participants encoded short news clips varying in emotional valence (Samide et al., 2020) and recalled them during fMRI scanning. Event memories persisting one day later were scored for narrative and perceptual details. We hypothesized that medial temporal subnetwork activity would be associated with perceptual detail memory, whereas dorsomedial and core midline subnetworks would relate to narrative detail memory. We further hypothesized that dorsomedial network activity would be modulated by emotional content, and that sensitivity to perceptual detail in the medial temporal network would be enhanced by negative valence.

In addition to univariate activity changes, emerging evidence indicates that activity patterns in DMN areas are sensitive to the specific details of events (Baldassano et al., 2017; Chen et al., 2017; Kuhl & Chun, 2014), and that pattern reinstatement during recall scales with how many details are remembered (Bird et al., 2015; Oedekoven et al., 2017). Less is known, however, about how these patterns correspond with the type of information remembered. One possibility is that event-specific recall patterns are influenced by narrative details that provide a consistent event model; alternatively, they might reflect recapitulation of perceptual details. To test these possibilities, we applied multivariate pattern analyses to examine whether DMN subnetworks represented event-specific details, and whether the strength and stability of these representations related to the persistence of narrative and perceptual details in memory.

## Methods

### Participants

This study recruited participants (N = 35, 20 female, 15 male) from the Greater Boston area, with an age range of 18-35 (M = 22.29). Participants were right-handed, spoke English fluently, and indicated no history of psychiatric or neurological conditions and no use of psychoactive medications. Two subjects were excluded due to excessive head motion, two subjects were excluded because they exited the scanner early due to discomfort, and one subject was excluded due to a computer error during their session, leading to a final sample of 30 (16 female, 14 male) participants. All participants provided informed consent as approved by the Boston College Institutional Review Board, and completed MRI and COVID safety screenings in compliance with Boston University Cognitive Neuroimaging Center protocols.

### Design

**Figure 1.**
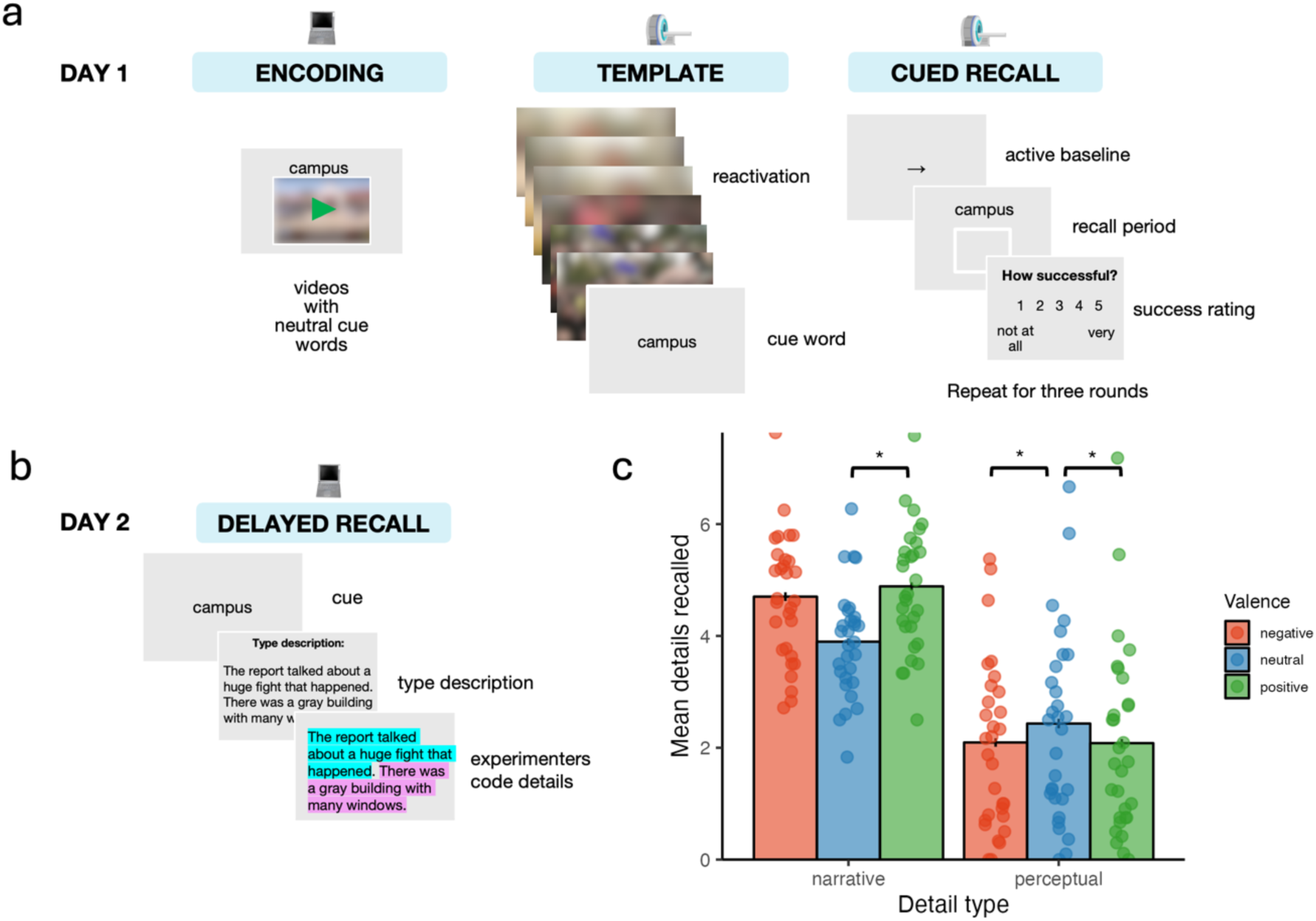
Experimental paradigm. **(a)** On experiment day 1, participants completed encoding outside of the scanner, followed by template and cued recall in an MRI scanner. Video images have been blurred in this figure for copyright reasons and bioRxiv policies. **(b)** On day 2, participants completed delayed recall outside of the scanner. **(c)** Average number of narrative and perceptual details recalled per video by valence category. More narrative details were remembered for positive than neutral events, while more perceptual details were remembered for neutral than positive or negative events. * indicates *p* < .05. Error bars represent standard error of the mean. Points reflect average values for each participant.

### Encoding Task

Before entering the scanner, participants first watched 48 clips of real-life news broadcasts of varying emotional valences (24 negative, 12 positive, 12 neutral) from the Samide et al. (2020) stimulus set (Figure 1a). Video clips were randomly presented, ranged from 20-52 seconds in length, and lasted 42 seconds on average. Each video showed a news report focused on one particular topic, such as a soldier homecoming, a weather report, or a murder. During each video, a semantically related, affectively neutral cue word was simultaneously displayed on the screen. For example, a video about a school chemistry lab fire used the cue word “chemistry”. Participants were informed they would be asked to recall the video from the cue word later. After each video, participants rated the valence of the clip, i.e., how emotionally negative or positive they found the clip, on a 9-point scale from 1 (most negative) to 9 (most positive) using buttons in the scanner. They then rated the intensity of the clip, i.e., how “emotionally intense” they found the clip, from 1 (least intense) to 9 (most intense). Participants were set up in the scanner immediately following the encoding phase.

### Template Task

In the scanner, participants first completed a “template” task. The purpose of this task was two-fold: one, to reinforce the video-cue associations after the participants were in the scanner, and two, to provide a “template” of brain activity corresponding to each video’s specific details, for comparisons with the later cued recall phase. Participants viewed a series of 10 chronological still images from each video (500 ms each × 10 = 5 second video period), with the relevant cue word displayed for 2 seconds after each video. In between trials, participants completed an active intertrial interval task (ITI) between each video in which they selected whether arrows appearing on the screen were pointing left or right. Each ITI period lasted eleven seconds. All video sets were shown once, divided across two functional runs.

### Cued Recall Task

The cued recall task consisted of three periods for each trial: “reactivation” (5 seconds), “elaboration” (10 seconds), and “success rating” (2 seconds). Participants were shown the previously assigned word cues and were instructed to silently recall the memory of the video associated with each cue. Participants pushed a button once they initially remembered the correct association. The “reactivation” period lasted five seconds, regardless of when the button was pushed. After five seconds, a box appeared around the cue word to signal the 10-second “elaboration” period. In total, participants remembered each video for 15 seconds per trial. For 2 seconds at the end of each trial, participants were prompted to rate their success in recalling the memory using a scale ranging from 1 (not at all successful) to 5 (very successful). Between each video, participants completed the same ITI task as in the template phase, with a duration of ten seconds.

Participants were cued to remember 12 negative, 12 positive and 12 neutral videos for each round of the cued recall task. This cued recall task was repeated for a total of three rounds, with each cue being presented once per round. In other words, each video was recalled three times. Each round separated into three consecutive runs, resulting in a total of 9 runs. Note that, for each participant, an additional 12 negative videos were recalled under instructions to reappraise the video during the elaboration period; these videos and the corresponding trials are excluded from the current analyses.

### Delayed Recall Task

The next day, participants completed a memory test online, with experimenter instructions provided via Zoom call. Participants typed out their memories for each video associated with the corresponding cue word on the screen. Participants were instructed to provide as much detail as possible in describing their memory of the video, and to include everything that came to mind in their description, such as storyline, setting, time, visual descriptions, thoughts, and feelings. Following each recall response, participants rated the vividness of the memory (1: least vivid, 9: most vivid), the valence of the memory (1: most negative, 9: most positive), and the emotional intensity of the memory (1: least intense, 9: most intense). In cases where participants could not remember the original video that matched the cue word, they were instructed to write “forgot”, and enter “0” for the subsequent ratings. There was no time limit to complete each recall response. Each video was recalled once (48 total cues displayed).

### Scoring

Cued recall data were hand-scored by counting the occurrence of a detail belonging to one of the following categories from the Autobiographical Interview scoring protocol (Levine et al., 2002): event, visual, place, time, emotion, reaction, extra, and repeated, which were then further labeled as correct or incorrect. A detail was considered “correct” if it accurately referenced any part of the video. The three categories used in the following analyses are “event” (happenings, actions, or people that are involved in the storyline), “visual” (purely perceptual aspects not integral to the storyline), and “correct” (the total number of correct details reported from any category). All other scored categories did not yield high enough detail counts for analysis and thus were unsuitable for inclusion in the present study. Here, narrative details are operationalized as event details, and perceptual details are operationalized as visual details.

### MRI Data Acquisition

MRI data were collected using a 3T Siemens MAGNETOM Prisma MRI scanner with a 32-channel head coil at the Boston University Cognitive Neuroimaging Center. Structural MRI images were collected using a T1-weighted multi-echo MPRAGE protocol (van der Kouwe et al., 2008) (field of view = 256 mm, GRAPPA (iPAT) acceleration = 4 (Griswold et al., 2002), 1 mm isotropic voxels, 176 sagittal slices with interleaved acquisition, TR = 2530 ms, TE = 1.69/3.55/5.41/7.27 ms, flip angle = 7 degrees, anterior-to-posterior phase encoding). Functional images were acquired using a whole-brain multiband (Moeller et al., 2010) echo-planar imaging sequence (bandwidth = 1718, field of view = 208 mm, 2 mm isotropic voxels, 69 slices with interleaved acquisition, TR = 1500ms, TE = 28ms, flip angle = 75, anterior-to-posterior phase encoding with a simultaneous multi-slice factor = 3 for a total of 296 TRs per scan run. Field map scans were acquired at the beginning of the scan session, then again after the third cued recall run to correct EPI images for signal distortion.

### Preprocessing and Modeling of fMRI data

MRI data were converted to NIfTI format using dcm2nii (Li et al., 2016) and converted to Brain Imaging Data Structure (BIDS) format using custom scripts in MATLAB (The Mathworks Inc., 2019), verified using the BIDS validator (http://bids-standard.github.io/bids-validator/). MRIQC v0.15.1 (Esteban, Birman, et al., 2017) was used to assess data quality. Scan runs were excluded from analyses if >20% of time points exceeded a framewise displacement of 0.30 mm.

MRI data were preprocessed using fmriprep v1.5.2 with the default processing steps (Esteban, Blair, et al., 2017). The brain was extracted and segmented from T1w images, which were then normalized to MNI space. BOLD images were skull stripped, slice timing corrected, field map corrected, and registered to the T1w images. Data remained unsmoothed to minimize blurring of signals across region-of-interest (ROI) boundaries. Framewise displacement, six realignment parameters, and six aCompCor components derived from fmriprep were included as nuisance regressors in each model.

After preprocessing, data were run in a model containing only the nuisance regressors listed above using SPM12. The residuals from this model were saved out in order to obtain corrected time series data. Residual activity was averaged across timepoints within a 15 second recall trial (a combination of the 5-second reactivation and 10-second elaboration period, corrected for hemodynamic lag) to obtain individual voxel activity for each trial. Then, activity was averaged across all voxels in each parcel to obtain ROI activity for each trial. Trials were excluded from analysis if any ROI activity values were more than 4 standard deviations from the subject’s mean (following Cooper & Ritchey, 2020).

### ROI definition

Neuroimaging data were extracted from the 200-parcel, 17-network cortical parcellations using the Schaefer-Yeo atlas (Schaefer et al., 2018). We targeted three default subnetworks: the core midline, the dorsomedial subnetwork, and the medial temporal subnetwork. Here, the core midline is delineated as DMN-A in the Schaefer-Yeo parcellation (Schaefer et al., 2018) and includes components of the medial prefrontal and parietal regions (anterior angular gyrus, posterior cingulate cortex). The dorsomedial subnetwork is referred to as DMN-B, including parcels in the dorsal and ventral prefrontal cortex and lateral temporal regions. The medial temporal subnetwork refers to medial temporal regions delineated as DMN-C and includes parcels in the parahippocampal cortex, ventral posterior cingulate, and posterior angular gyrus. To verify that these groupings were meaningful in our dataset, we examined within- and between-network parcel correlations and found that parcels from the same subnetwork were coactivated more strongly with each other during the task than with parcels from other subnetworks (Supplementary Figure 1).

Four additional medial temporal lobe (MTL) regions of interest (ROIs) were added for left and right hippocampus and amygdala. The hippocampus was obtained from a probabilistic MTL atlas (Ritchey et al., 2015; https://neurovault.org/collections/3731/) and amygdala was added from the Harvard-Oxford subcortical atlas, in line with prior methods (Cooper & Ritchey, 2019). Six additional hippocampal subfield ROIs (left and right head, body, and tail) were also used for secondary analyses (Ritchey et al., 2015; https://neurovault.org/collections/3731/), with body and tail collapsed to represent the posterior hippocampus.

### Statistical Analysis

All statistical analyses were conducted using R 4.3.2 (https://www.r-project.org/). For the behavioral analyses, ANOVAs were used to examine the effect of emotional valence (positive, neutral, negative) on the average number of narrative, perceptual, and overall correct details recalled, using the R-package *ez* (Lawrence, 2016); posthoc comparisons were computed using paired-samples t-tests. For the fMRI analyses, trial-wise activity in each network was predicted using linear mixed-effects models, which included the subject as a random intercept utilizing the R-packages *lme4* (Bates et al., 2015) and *lmerTest* (Kuznetsova et al., 2017). We tested models including random slopes for valence and details; however, these models were often singular or failed to converge, and thus we report the random-intercept-only models here for consistency across the models. In order to maximize the variance accounted for by the model, all predictors were included in each model. Because all predictors were included simultaneously, regression coefficients reflect each predictor’s unique association with network activity, accounting for the other predictors. Analyses followed the subsequent equation:

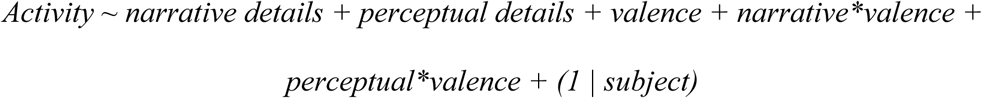

This model was fit individually for each network, where network activity corresponded to the average across ROIs within a network. Due to the categorical nature of “valence” (positive, neutral, negative), analyses were first performed using “neutral” as the reference category, then rerun using “positive” as the reference category to capture any additional differences between positive and negative valence that could not be tested using “neutral” as the reference category. For each effect, p-values were adjusted for multiple comparisons using the Benjamini-Hochberg method with a false discovery rate (FDR) of .05 (Benjamini & Hochberg, 1995), taking into account the tests for each network and MTL ROI. Values for narrative and perceptual details were mean centered within participants to better account for between-subjects variability. In order to examine the effect of narrative and perceptual details beyond a simple difference between remembered and forgotten videos (no details), our analyses included only trials classified as “remembered” as indicated by participant rating of recall success at delayed retrieval. We also explored whether these results held up when considering all trials (remembered and forgotten). Note that all reported results remained significant except for one finding noted in the Results section.

### Representational Similarity Analysis

In order to assess the reactivation of event-specific information during recall, a representational similarity analysis (RSA; Kriegeskorte et al., 2008) was conducted to compare multi-voxel pattern similarity between corresponding trials in the template and cued recall phases (Figure 4a). The spatial pattern evoked by each recall trial was compared to the pattern for each template phase trial using Pearson’s correlation, and resulting correlation values were Fisher z transformed. Event-specific reactivation was defined by the difference between same-video comparisons and different-video comparisons, computed separately for each recall trial. Same-video comparisons involved trial correlations that were matched on video, while different-video comparisons involved differing videos but were matched for valence and whether they were remembered or forgotten in the delayed recall task. Note that this was computed separately for each of the 3 recall trials associated with each video. This measure was evaluated separately for each parcel and then combined for the 3 default mode subnetworks. Trial-wise network values were entered into a regression model similar to the one used for the univariate analyses, with the z-transformed difference between same and different video trials included as the outcome variable. Each effect was adjusted for multiple comparisons using the Benjamini-Hochberg method with a FDR of .05.

## Results

### Valence effects on memory details

One day after encoding news videos and covertly retrieving them in the MRI, participants were asked to overtly recall as many details as possible about each video. To evaluate whether these memory measures at delayed recall related to in-scanner behavioral measures at cued recall, we related these details to participants’ success ratings collected after each trial of cued recall in the MRI, while accounting for the effects of emotional valence. Mean success ratings significantly predicted delayed vividness ratings (*β* = 0.720, *t*(989) = 5.161, *p* < .001) as well as the number of perceptual and narrative details retained after one day (perceptual: *β* = 0.341, *t*(989) = 2.163, *p* < .05; narrative: *β* = 0.439, *t*(989) = 2.827, *p* < .01), indicating that Day 1 subjective memory success relates to the strength of memory on Day 2. Participants generally reported high levels of success among trials that were successfully recalled the following day (i.e., those included in the present analyses): 86.8% of trials received a rating of 4 or 5 on the 1-5 success scale (Supplementary Figure 2). These findings indicate that participants were generally successful in bringing video details to mind during the scan, and that their in-scanner recall was likely to correspond with the details recalled in the delayed test.

We then examined the effect of video valence (positive, neutral, negative) on the average number of overall, narrative, and perceptual details recalled by means of an ANOVA. There was a significant effect of valence on overall correct details remembered, *F*(2,58) = 4.32, *p* < .05, η²_G_ = .014, indicating that participants recalled more correct details for positive videos (*M* = 7.95, *SD* = 3.95) than for neutral (*M* = 7.18, *SD* = 3.47, *t*(29) = 3.028, *p* < .01).

We then distinguished between narrative and perceptual details to examine any distinct effects within detail types. Here, we found a significant effect of valence, *F*(2,58) = 22.64, *p* < .001, η²_G_ = .13 (Figure 1b), such that more narrative details were recalled from positive videos (*M* = 4.89, *SD* = 2.21, *t*(29) = 6.063, *p* < .001) and negative videos (M = 4.70, *SD* = 2.31, *t*(29) = 5.404, *p* < .001) than neutral (*M* = 3.90, *SD* = 1.87). For perceptual details, on the other hand, there was a memory benefit for neutral videos, *F*(2,58) = 3.87, *p* < .01, η²_G_ = .009 (Figure 1b), such that participants recalled more perceptual details for neutral videos (*M* = 2.43, *SD* = 2.68) compared to negative (*M* = 2.09, *SD* = 2.26, *t*(29) = -2.79, *p* < .01) or positive videos (*M* = 2.08, *SD* = 2.32, *t*(29) = 2.509, *p* < .05). Thus, our results indicate differential effects of valence on memory for narrative versus perceptual details, in which narrative details were remembered best for positive and negative videos whereas perceptual details were recalled best for neutral videos.

Importantly, narrative details and perceptual details were only moderately correlated with each other across participants (*r* = 0.376, *p* < .05), suggesting that there may be separate mechanisms by which narrative and perceptual details contribute to total remembering.

### Activity related to memory for narrative and perceptual details

In order to examine how default mode subnetwork activity relates to the contents of later memory and the valence of the remembered event, activity was first averaged across ROIs for each of the dorsomedial, medial temporal, and core midline subnetworks. Mixed-effects linear regression analyses were used to predict trial activity as a function of valence and the number of recalled narrative and perceptual details.

We found that activity in the dorsomedial subnetwork was sensitive to negative valence, such that negative videos produced higher activity in the dorsomedial network than for neutral videos (*β* = .065, CI: .03 - .1, *t*(2724.064) = 3.678, *p_FDR_* = .001, Figure 2a, c). Furthermore, we found that the core midline activity was significantly predicted by positive valence relative to neutral (*β* = .041, CI: .01 - .07, *t*(2722.820) = 2.964, *p_FDR_* = .015). Note that this latter finding was specific to analyses limited to remembered trials and did not emerge when combining remember and forgotten trials together. Core midline and dorsomedial subnetwork activity was not related to memory for narrative or perceptual details. In contrast, medial temporal subnetwork activity scaled with the number of perceptual details recalled (*β* = .011, CI: .00 - .02, *t*(2726.857) = 2.509, *p_FDR_* = .042, Figure 2d), regardless of the emotional valence of the memory. When the reference category was switched to positive to assess positive-negative comparisons, no additional significant comparisons emerged, *p_FDR_* > .05, and the relationship with perceptual details became non-significant, likely due to a numerically weaker relationship for the positive reference category. Providing further support for the relationship with perceptual details recalled, we also found that medial temporal subnetwork activity was positively associated with success ratings during in-scanner recall (Supplementary Figure 2).

**Figure 2.**
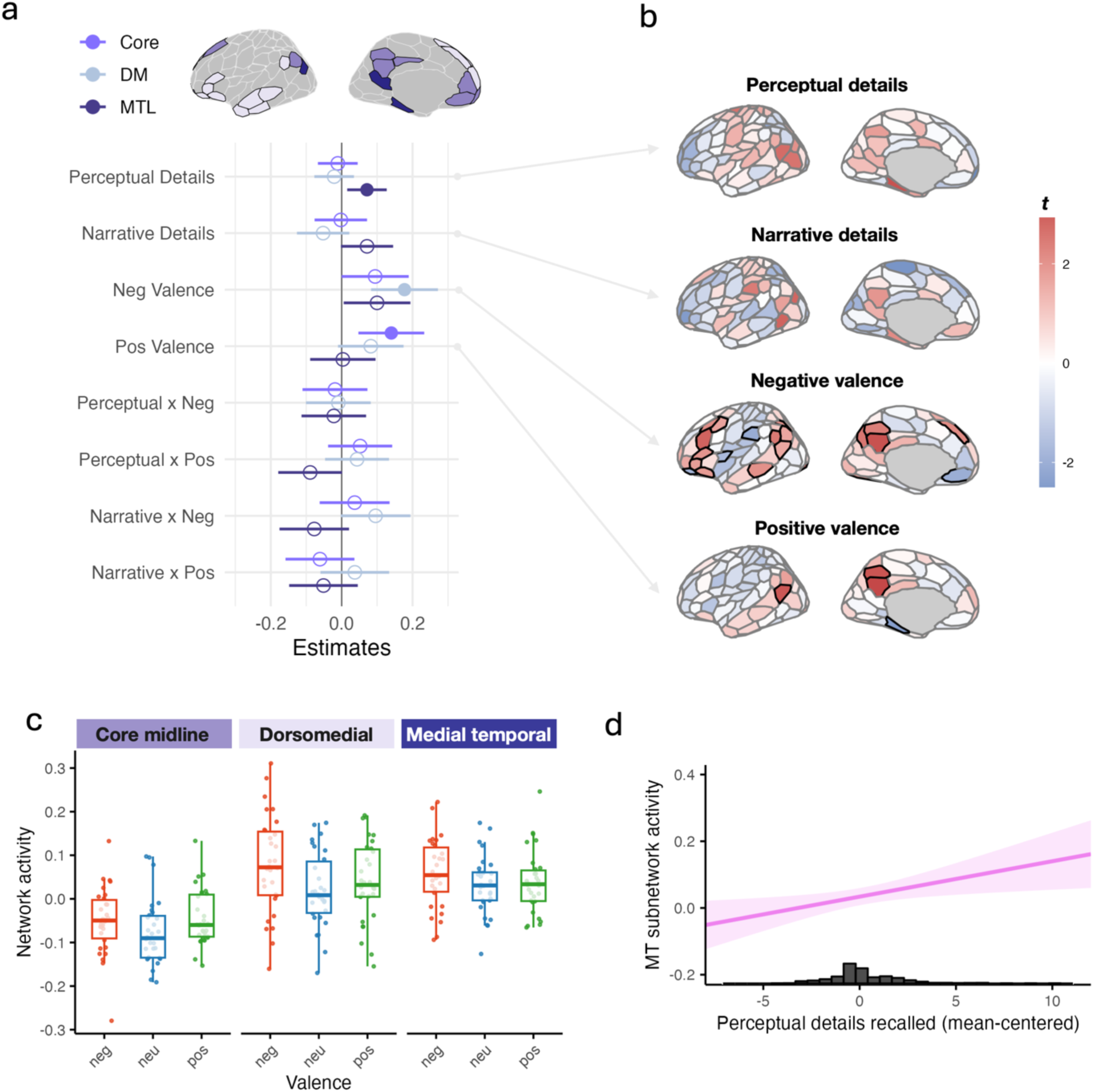
DMN activity related to remembered event features. **a)** Mixed-effects model regression estimates for each DMN subnetwork with network activity as the outcome variable. Error bars denote the 95% confidence interval for each estimate. Filled circles denote FDR-corrected *p* < .05. Networks are displayed using the Schaefer parcellation (Schaefer et al., 2018) with the Yeo 17-network labels (Yeo et al., 2011). **b)** Whole-brain mixed-effects model results for the four main predictors, displayed with the Schaefer parcellation (Schaefer et al., 2018). Parcels with significant t-values after FDR correction are outlined in black. **c)** DMN subnetwork activity by valence. Boxplots represent the distribution of average network activity for each valence category, with the boxplots showing the median and the interquartile range across participants and with individual data points representing each participant. **d)** Medial temporal subnetwork activity scales with the number of perceptual details recalled. Shading denotes the 95% confidence interval around the predicted values. The mixed-effects model regressor corresponding to this comparison was significant at FDR-corrected *p* < .05. Histogram represents the mean-centered distribution of perceptual details recalled, including trials from all participants.

To characterize these effects further and to examine the involvement of individual parcels in our network-level effects, we ran the same models for all 200 parcels in the brain. Here, we focus on the results for DMN parcels (Table 1); results for non-DMN parcels are reported in Figure 2b and Supplementary Table 1. We found that valence-related effects were distributed but most prevalent within the core and dorsomedial subnetworks (Table 1, Supplementary Figure 3), consistent with our network-level findings. No individual DMN parcel showed significant sensitivity to narrative or perceptual detail. However, visual inspection of parcel-level effects suggests that the medial temporal subnetwork shows coherent recruitment specifically for detail-related effects, suggesting that subnetwork-level specialization may emerge from distributed activity rather than isolated parcel effects (Supplementary Figure 3).

**Table 1.**
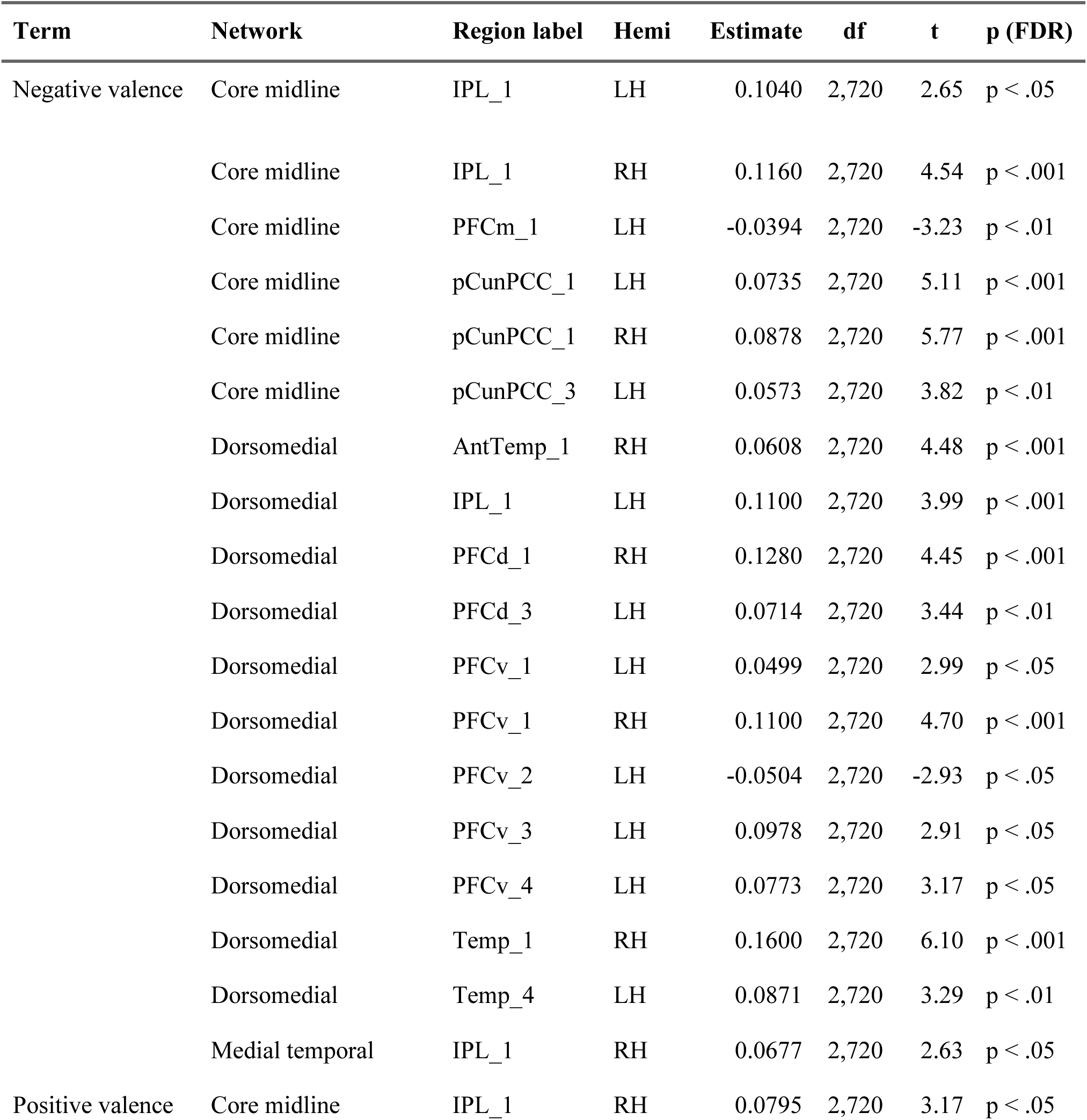
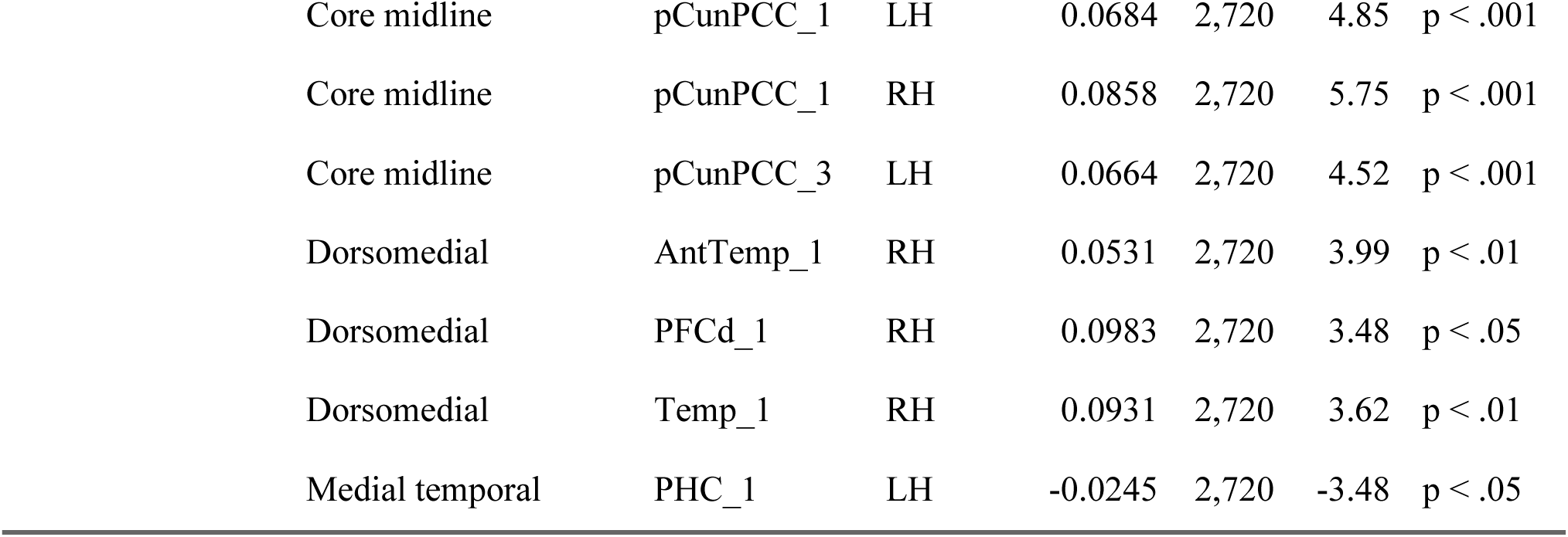
Significant parcel-level outcomes across all parcels in the DMN for the main mixed-effects model estimates, where FDR-corrected *p* < .05. Region labels are from the Schaefer atlas. LH=left hemisphere, RH=right hemisphere.

| Term | Network | Region label | Hemi | Estimate | df | t | p (FDR) |
| --- | --- | --- | --- | --- | --- | --- | --- |
| Negative valence | Core midline | IPL_1 | LH | 0.1040 | 2,720 | 2.65 | $p < .05$ |
| | Core midline | IPL_1 | RH | 0.1160 | 2,720 | 4.54 | $p < .001$ |
| | Core midline | PFCm_1 | LH | -0.0394 | 2,720 | -3.23 | $p < .01$ |
| | Core midline | pCunPCC_1 | LH | 0.0735 | 2,720 | 5.11 | $p < .001$ |
| | Core midline | pCunPCC_1 | RH | 0.0878 | 2,720 | 5.77 | $p < .001$ |
| | Core midline | pCunPCC_3 | LH | 0.0573 | 2,720 | 3.82 | $p < .01$ |
| | Dorsomedial | AntTemp_1 | RH | 0.0608 | 2,720 | 4.48 | $p < .001$ |
| | Dorsomedial | IPL_1 | LH | 0.1100 | 2,720 | 3.99 | $p < .001$ |
| | Dorsomedial | PFCd_1 | RH | 0.1280 | 2,720 | 4.45 | $p < .001$ |
| | Dorsomedial | PFCd_3 | LH | 0.0714 | 2,720 | 3.44 | $p < .01$ |
| | Dorsomedial | PFCv_1 | LH | 0.0499 | 2,720 | 2.99 | $p < .05$ |
| | Dorsomedial | PFCv_1 | RH | 0.1100 | 2,720 | 4.70 | $p < .001$ |
| | Dorsomedial | PFCv_2 | LH | -0.0504 | 2,720 | -2.93 | $p < .05$ |
| | Dorsomedial | PFCv_3 | LH | 0.0978 | 2,720 | 2.91 | $p < .05$ |
| | Dorsomedial | PFCv_4 | LH | 0.0773 | 2,720 | 3.17 | $p < .05$ |
| | Dorsomedial | Temp_1 | RH | 0.1600 | 2,720 | 6.10 | $p < .001$ |
| | Dorsomedial | Temp_4 | LH | 0.0871 | 2,720 | 3.29 | $p < .01$ |
| | Medial temporal | IPL_1 | RH | 0.0677 | 2,720 | 2.63 | $p < .05$ |
| Positive valence | Core midline | IPL_1 | RH | 0.0795 | 2,720 | 3.17 | $p < .05$ |

**Table 1.** Significant parcel-level outcomes across all parcels in the DMN for the main mixed-effects model estimates, where FDR-corrected $p < .05$ . Region labels are from the Schaefer atlas.
|  |  |  |  |  |  |  |
| --- | --- | --- | --- | --- | --- | --- |
| Core midline | pCunPCC_1 | LH | 0.0684 | 2,720 | 4.85 | $p < .001$ |
| Core midline | pCunPCC_1 | RH | 0.0858 | 2,720 | 5.75 | $p < .001$ |
| Core midline | pCunPCC_3 | LH | 0.0664 | 2,720 | 4.52 | $p < .001$ |
| Dorsomedial | AntTemp_1 | RH | 0.0531 | 2,720 | 3.99 | $p < .01$ |
| Dorsomedial | PFCd_1 | RH | 0.0983 | 2,720 | 3.48 | $p < .05$ |
| Dorsomedial | Temp_1 | RH | 0.0931 | 2,720 | 3.62 | $p < .01$ |
| Medial temporal | PHC_1 | LH | -0.0245 | 2,720 | -3.48 | $p < .05$ |

Similar analyses were conducted for the hippocampal and amygdala ROIs, motivated by the known role of the hippocampus and amygdala in emotional memory (Phelps, 2004) and potential dissociations among hippocampal subregions in emotional memory (Ritchey et al., 2019). Overall hippocampal activity was significantly predicted by the number of remembered perceptual details (*β* = .005, CI: .00 - .01, *t*(2728.025) = 2.192, *p_FDR_* = .047). A hippocampal subregion analysis revealed that this relationship was significant for the left anterior hippocampus (*β* = .008, CI: .00 - .01, *t*(2722.682) = 2.544, *p_FDR_* = .044; Figure 3) but not for the hippocampal body or tail. Amygdala activity also significantly scaled with the number of remembered perceptual details (*β* = .007, CI: .00 - .01, *t*(2726.404) = 2.389, *p_FDR_* = .042; Figure 3), and in the models directly comparing positive and negative valence, amygdala activity was significantly predicted by positive valence compared to negative (*β* = -.028, CI: -.05 - -.01, *t*(2721.456) = -2.968, *p_FDR_* = .003). Neither hippocampal nor amygdala activity was related to the number of narrative details recalled. All other effects across models and ROIs were non-significant, all *p_FDR_* > .05.

**Figure 3.**
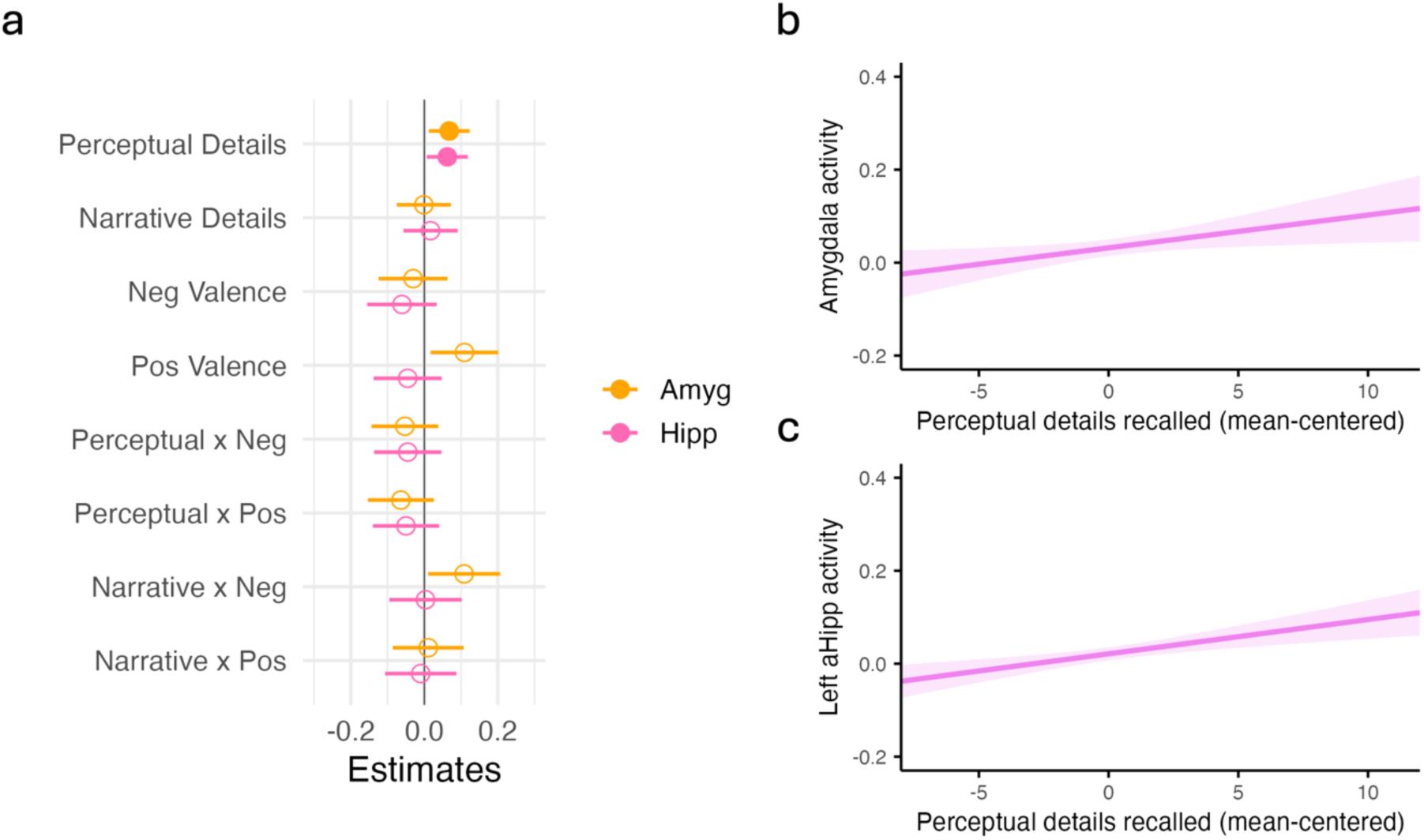
Amygdala and hippocampal activity related to remembered event features. **a)** Mixed-effects model estimates for amygdala and hippocampus with activity as the outcome variable. Error bars denote the 95% confidence interval for each estimate. Filled circles denote FDR-corrected *p* < .05. **b)** Amygdala and **c)** left anterior hippocampus activity scale with the amount of perceptual details recalled. Shading denotes the 95% confidence interval around the predicted values. The mixed-effects model regressor corresponding to this comparison was significant at FDR-corrected *p* < .05.

Together, these results suggest that the dorsomedial and core midline subnetworks are especially responsive to memories for emotional events, whereas the medial temporal subnetwork, hippocampus, and amygdala are more active for memories with more perceptual details.

### Event-specific patterns of activation and their relation to memory quality

To determine how the strength of event-specific patterns in the DMN is related to recapitulation of different types of details, we conducted a representational similarity analysis (RSA) assessing multi-voxel pattern similarity between the template and cued recall phases of the experiment, and its relationship to detail type and emotional valence. We reasoned that this measure of template-recall similarity would be sensitive to the reactivation of high-fidelity, event-specific details from encoding through the recall phase. Across conditions and in each of the DMN subnetworks, there was evidence for event-specific reactivation (Figure 4b), such that same-video similarity was greater than different-video similarity (core midline: *t*(29) = 3.94, dorsomedial: *t*(29) = 3.52, medial temporal: *t*(29) = 5.13; all *p* < .001). However, this effect was not significantly associated with the number of narrative or perceptual details or with negative or positive valence compared against neutral, *p_FDR_* > .05. When comparing negative and positive valence directly (i.e., with positive valence as the reference), there was an interaction between valence and narrative details, such that reactivation of medial temporal subnetwork areas was more strongly associated with narrative detail for negative videos compared to positive (*β* = .005, CI: .00 - .01, *t*(2788.307) = 3.03, *p_FDR_* = .007, Figure 4c). Finally, we note that there was a significant relationship with perceptual details in the positive reference model that was not observed in the neutral reference model, likely due to numerical differences in the strength of the relationship for the different valence categories. Thus, to clarify these effects, we additionally ran a simpler model collapsing across valence, including fixed effects only for narrative and perceptual details, and found that there was no significant relationship between reactivation in DMN areas and memory for either kind of detail, *p_FDR_* > .05. These findings suggest that reactivation in the medial temporal network may be more influenced by narrative detail for negative compared to positive events, but that otherwise, there was little association between template-recall similarity and memory quality.

**Figure 4.**
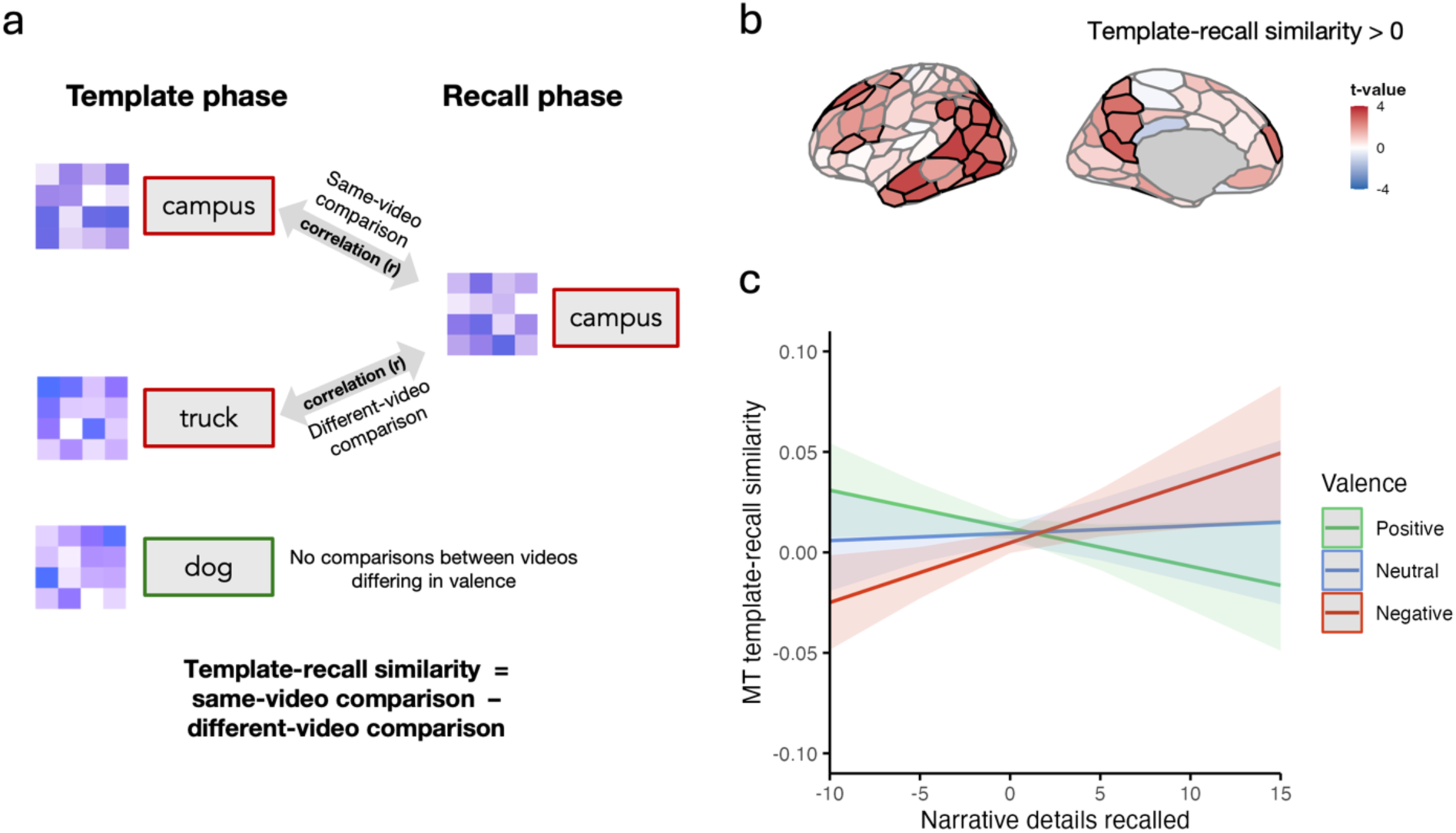
Template-recall similarity. **a)** Template-recall similarity RSA method. Same-video comparisons were calculated by correlating activity patterns elicited by the same video during the template and recall phases. Different-video comparisons were calculated by correlating each trial’s recall activity pattern with the template activity pattern of other videos matched on valence. **b)** Whole-brain results of a one-sample t-test of template-recall similarity (same-video similarity minus different-video similarity) displayed with the Schaefer parcellation (Schaefer et al., 2018). Parcels with significant t-values after FDR correction are outlined in black. **c)** Template-recall similarity in the medial temporal subnetwork was more strongly positively related to narrative details for negative compared to positive valenced events. Shading denotes the 95% confidence interval around the predicted values. The mixed-effects model interaction term corresponding to this comparison was significant at FDR-corrected *p* < .05.

In order to capture the consistency, or stability, of event-specific memory representations across recalls, another RSA was conducted to examine multivoxel pattern similarity across the three recall rounds in the cued recall task (Figure 5a). The spatial pattern of activity for each recall-phase trial was correlated with that of every other recall-phase trial using Pearson’s correlation, and resulting correlation values were Fisher z transformed. For each trial, same-video comparisons were made with patterns corresponding to the same video across the other rounds, and different-video comparisons were made with different videos from the other rounds that were matched for valence and whether they were remembered or forgotten in the delayed recall task. Recall stability was calculated as the difference between same-video comparisons and different-video comparisons, evaluated separately for each parcel and then combined for the 3 default mode subnetworks. Trial-wise values were saved out for each ROI using the same atlas and ROIs as in the univariate analysis described previously. The primary analysis used a mixed-effects regression model which included narrative and perceptual details as predictors, and a secondary analysis included emotional valence as an additional predictor to match the regression model described above. The z-transformed difference between same and different video trials was included as the outcome variable. This model was run for each of the 3 default mode subnetworks, and each effect was adjusted for multiple comparisons using the Benjamini-Hochberg method with a FDR of .05 (Benjamini & Hochberg, 1995). For both the template-recall and recall-recall approaches, one-sample t-tests were computed to test whether event-specific reactivation (averaged across trials for each participant) in each of the DMN subnetworks was greater than zero.

Because event patterns may be transformed from perception to recall (Favila et al., 2020; Xiao et al., 2017; Xue, 2022), we additionally assessed how event-specific memory representations were maintained across the three retrieval rounds in the cued recall phase. We reasoned that this measure of recall-recall similarity would indicate the stability of the event representation over repeated retrievals. Across conditions and in each of the DMN subnetworks, there was evidence for event-specific recall stability (Figure 5b), such that same-video similarity was greater than different-video similarity (core midline: *t*(29) = 5.13, dorsomedial: *t*(29) = 6.09, medial temporal: *t*(29) = 4.60; *p* < .002). In addition, recall stability was significantly modulated by the contents of memory. Recalling positively valenced events was associated with more stable recall patterns in core midline and dorsomedial subnetworks than recalling neutral events (core midline: *β* = .009, CI: .00 - 02, *t*(2763.135) = 2.813, *p_FDR_* = .009; dorsomedial: *β* = .012, CI: .00 - .02, *t*(2765.065) = 3.213, *p_FDR_* = .004; Figure 5c). Recall stability was also stronger in core midline for negative event recall compared to neutral (*β* = .009, CI: .00 - 02, *t*(2764.231) = 2.745, *p_FDR_* = .028, Figure 5c). The simpler model containing only narrative and perceptual details as predictors showed that perceptual detail recall scaled with recall stability in the medial temporal network (*β* = .002, CI: .00 - .00, *t*(2763.168) = 3.34, *p_FDR_* = .003). All other effects across models and networks were non-significant, all *p_FDR_* > .05. These results suggest that event-specific representations of valence memory are maintained across retrievals in the dorsomedial and core midline subnetworks, whereas representations in the medial temporal network are sensitive to perceptual details independently of valence.

**Figure 5.**
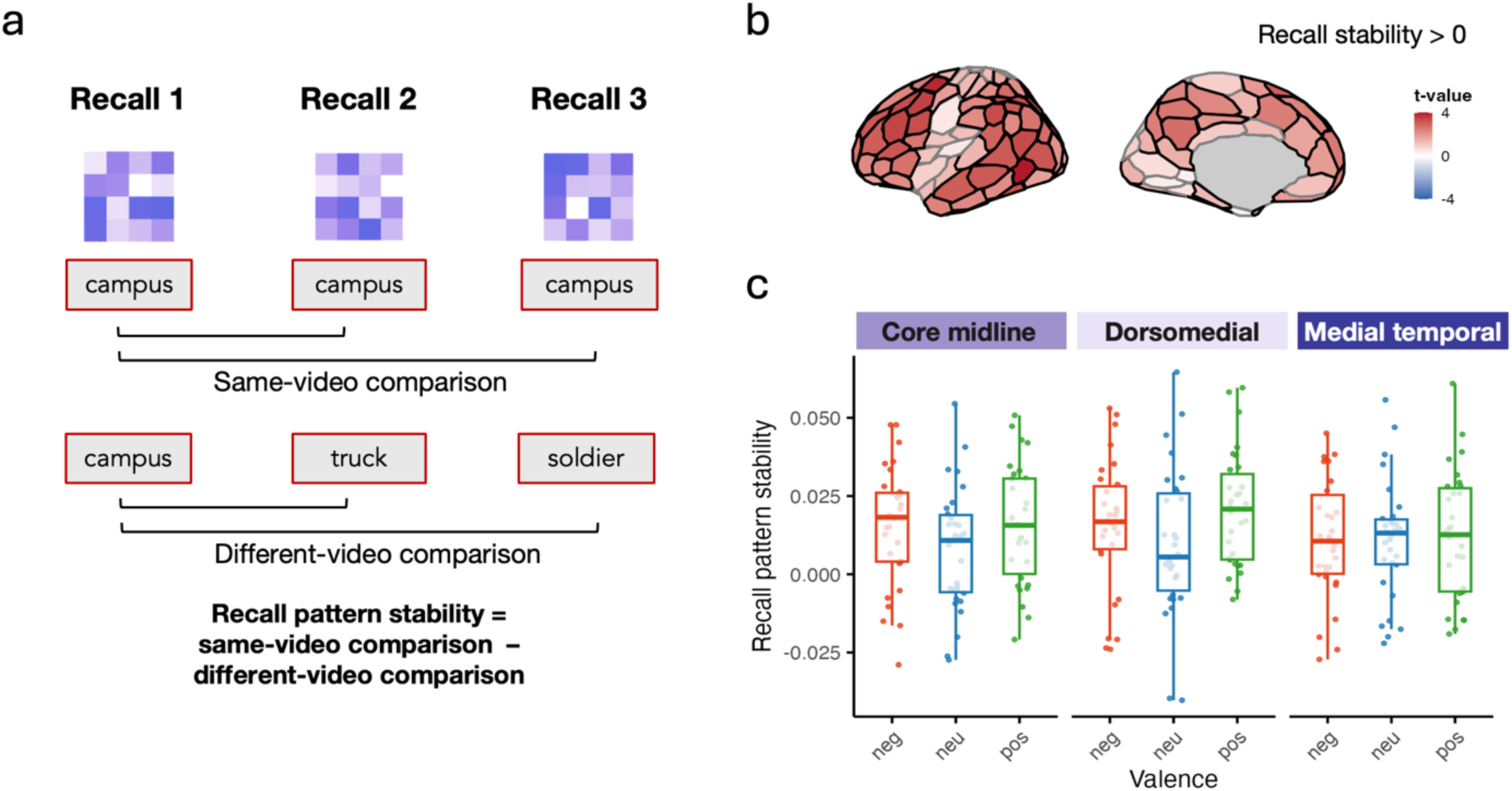
Recall stability. **a)** Recall stability RSA method. Each trial’s retrieval pattern was compared to other repetitions of the same video recall (i.e., in different rounds) to obtain the average same-video comparison. The trial’s correlations with other valence-matched, different-video recall trials were averaged to obtain the different-video comparison. **b)** Whole-brain results of a one-sample t-test of recall pattern stability (same-video comparisons minus different-video comparisons) displayed with the Schaefer parcellation (Schaefer et al., 2018). Parcels with significant t-values after FDR correction are outlined in black. **c)** DMN subnetwork recall stability by valence. Boxplots represent the distribution of average network recall stability for each valence category, with the boxplots showing the median and the interquartile range across participants and with individual data points representing each participant.

To examine the contribution of individual parcels to these results, we re-ran both our RSA approaches across all 200 parcels in the brain. We found that parcels in the dorsomedial and core networks showed greater recall stability for positive memories compared to neutral memories, with a similar trend for negative compared to neutral memories (Supplementary Figure 4). One region in the dorsomedial network, the ventromedial prefrontal cortex, was also sensitive to perceptual details recalled. For template-recall similarity, only one DMN parcel showed evidence of being modulated by the features of the remembered videos (Supplementary Figure 5): the right parahippocampal cortex showed reactivation more for positive than neutral videos, with a similar non-significant pattern for negative videos (Table 2). Whole-brain results outside of the DMN can be found in Supplementary Table 2. These results indicate that recall stability in parcels across the dorsomedial and core networks is generally modulated by valence, but that the relationship with details may be more variable across parcels.

**Table 2.**
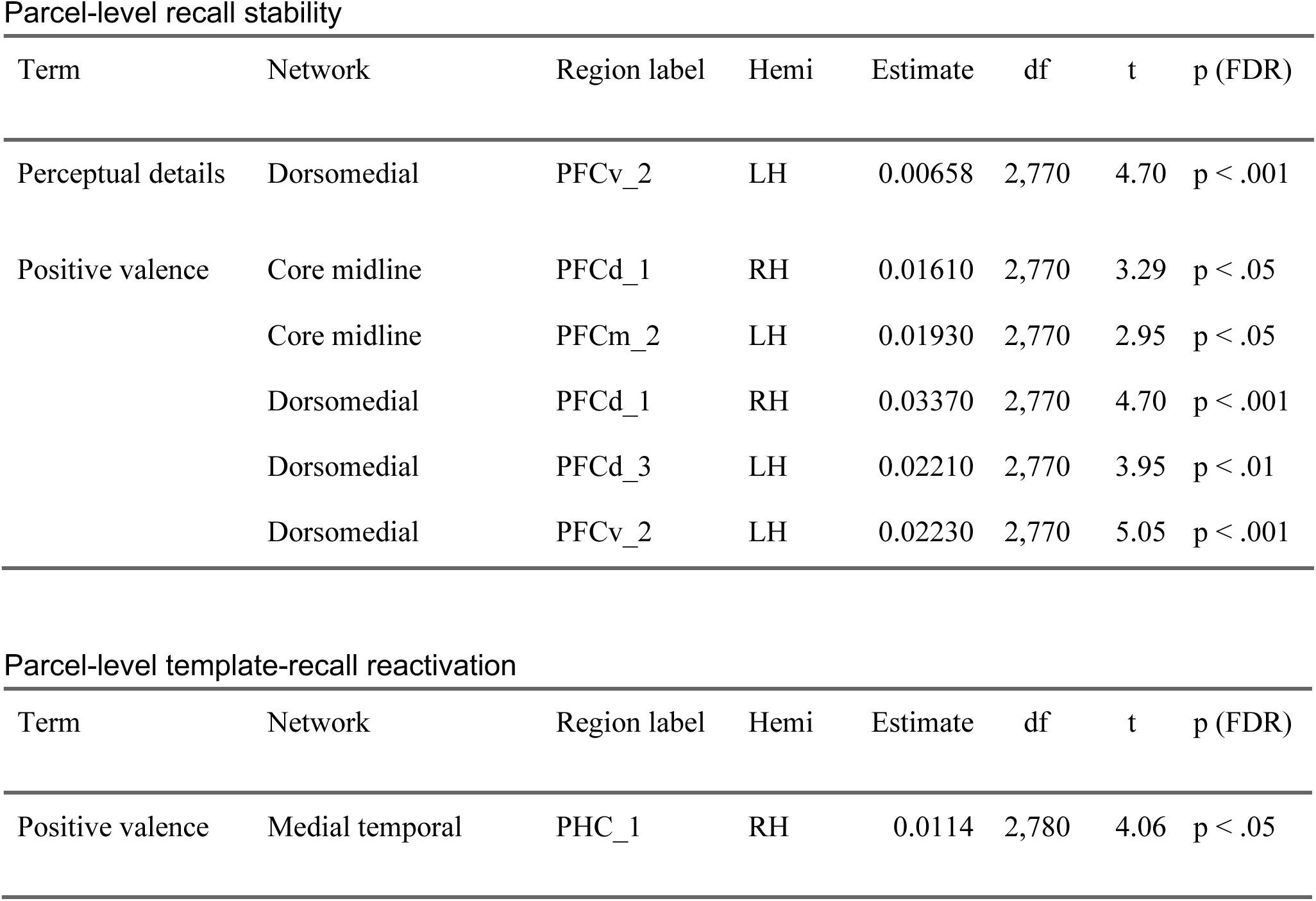
Significant parcel-level outcomes across all parcels in the DMN for the mixed-effects model estimates from each RSA approach, where FDR-corrected *p* < .05. Region labels are from the Schaefer atlas. LH=left hemisphere, RH=right hemisphere.

Parcel-level recall stability
| Term | Network | Region label | Hemi | Estimate | df | t | p (FDR) |
| --- | --- | --- | --- | --- | --- | --- | --- |
| Perceptual details | Dorsomedial | PFCv_2 | LH | 0.00658 | 2,770 | 4.70 | $p < .001$ |
| Positive valence | Core midline | PFCd_1 | RH | 0.01610 | 2,770 | 3.29 | $p < .05$ |
| | Core midline | PFCm_2 | LH | 0.01930 | 2,770 | 2.95 | $p < .05$ |
| | Dorsomedial | PFCd_1 | RH | 0.03370 | 2,770 | 4.70 | $p < .001$ |
| | Dorsomedial | PFCd_3 | LH | 0.02210 | 2,770 | 3.95 | $p < .01$ |
| | Dorsomedial | PFCv_2 | LH | 0.02230 | 2,770 | 5.05 | $p < .001$ |

## Discussion

The present study examined whether default mode network (DMN) contributions to recall vary by the valence and detailedness of memories. We found that the engagement of DMN subnetworks varied depending on the features persisting in memory: dorsomedial subnetwork activity was associated with memory for event valence, while medial temporal subnetwork activity scaled with the number of perceptual details recalled. Amygdala and anterior hippocampus activity also scaled with remembered perceptual details, with the amygdala also showing greater activity for positive versus negative events. In addition to these activation differences, event-specific patterns in the DMN, and in particular, their stability across recall repetitions, were also affected by memory valence and detail. These findings demonstrate that DMN contributions to recall are modulated by emotional valence and the persistence of specific details in memory.

Our finding that medial temporal subnetwork activity was related to the perceptual detailedness of memory aligns with previous studies implicating these regions in episodic retrieval. The medial temporal subsystem is involved in episodic memory and scene construction (Andrews-Hanna et al., 2010) and is more strongly connected to visual network regions than other DMN components (Barnett et al., 2021). Here, the relationship with perceptual details emerged at the network level: each parcel within the medial temporal subnetwork showed a similar profile of results, suggesting that the network-level effect was not carried by any individual region but rather the network as a whole. Recall stability in the medial temporal subnetwork was also related to the amount of remembered perceptual details, suggesting that event-specific patterns in these areas are intertwined with perceptual aspects of memory. Contrary to our hypothesis, this relationship was not modulated by negative emotional valence, despite prior findings that negative valence biases visual processing (Kark & Kensinger, 2019) and enhances sensory recapitulation (Bowen et al., 2018). Behaviorally, neutral events were remembered with more perceptual detail than emotional events. This discrepancy may stem from the nature of our naturalistic news stimuli, where the central details were largely narrative aspects of the event, rather than vividly perceptual in nature (as compared with studies using threatening or graphic images). Interestingly, however, valence did modulate the relationship between narrative detail and template-recall similarity in the medial temporal subnetwork, with a stronger relationship for negative compared to positive stimuli. This unexpected finding suggests that high-fidelity recapitulation may serve to protect the narrative of memory more strongly for naturalistic negative events.

Additional analyses implicated the amygdala and hippocampus in perceptual recall, with left anterior hippocampus activity scaling positively with recalled perceptual details. This is consistent with studies linking anterior hippocampus to greater pattern specificity (Ezzyat et al., 2018; Tompary & Davachi, 2017), scene construction (Zeidman & Maguire, 2016), and detailed episodic processes (Addis et al., 2011; Dandolo & Schwabe, 2018). Posterior hippocampus activity was unrelated to any of our predictors, and narrative detail did not predict anterior or posterior hippocampus activity. Thus, in contrast with some prior proposals (Poppenk et al., 2013; Robin & Moscovitch, 2017; Sheldon et al., 2019), we found no evidence for differential sensitivity to narrative versus perceptual recall along the hippocampal long axis.

Our results support the hypothesis that the dorsomedial network represents the emotional valence of memory. Dorsomedial DMN subdivisions have been linked to emotional experience (Andrews-Hanna et al., 2010; Barnett et al., 2021), with the dmPFC as a key node (Kensinger & Ford, 2021). Because memories were recalled in response to neutral cue words, our findings reflect valence information arising from memory rather than external cues. The dorsomedial subnetwork exhibited greater recall stability for positive versus neutral events, indicating that it maintains particularly consistent representations of positively valenced individual events (regardless of the number of details recalled). This subnetwork may be sensitive to emotional information in general rather than being valence-specific, as there were no significant differences in activity or recall stability when directly comparing positive and negative valence. Parcel-level analyses showed widespread activation of dorsomedial DMN regions associated with memory valence, though there was some variability across parcels within the subnetwork (Supplementary Figure 3). Contrary to our original hypothesis, the dorsomedial subnetwork did not show a stronger association with narrative details for emotional memories. This, along with its involvement in emotional recall regardless of the number of remembered details, suggests that the dorsomedial subnetwork primarily provides an affective scaffold for memory, integrating emotional tone without directly calling upon specific narrative or perceptual components.

The core midline subnetwork showed a similar profile of activity to the dorsomedial subnetwork, but with some differences related to emotional valence. Specifically, it was more active for positive than neutral events,and exhibited more stable event-specific retrieval patterns for both negative and positive events compared to neutral. These results suggest a role in integrating affective experience into memory, consistent with evidence that core midline regions activate together during self-relevant affective decisions (Andrews-Hanna et al., 2010). These results point to similarities in dorsomedial and core midline contributions to emotional memory recall, consistent with recent proposals that these are not entirely distinct networks. Individual-level functional connectivity analyses have demonstrated that the core midline’s prominence in group-averaged data results from anatomical interdigitation between the two primary networks identifiable at the individual level (Braga et al., 2019), and studies remain agnostic about a clear function for this region (Wen et al., 2020). Thus, the core midline’s sensitivity to affective valence in this study may result from signal intermixing from the dorsomedial subnetwork.

Our findings contribute to existing evidence of functionally distinct DMN subnetworks supporting episodic memory and cognition (Andrews-Hanna et al., 2010; Barnett et al., 2021; Ranganath & Ritchey, 2012). Prior research suggests that dorsal DMN supports abstract representations while ventral DMN supports perceptually specific information (Andrews-Hanna & Grilli, 2021; Ritchey & Cooper, 2020; Sheldon et al., 2019; Spunt et al., 2016). Furthermore, dorsal and ventral DMN are modulated by the emotional valence and vividness, respectively, of imagined future events (Lee et al., 2021), and are differentially recruited when generating emotional versus semantic contextual associations (Souter et al., 2024). Here, we find that such differences extend to contents of episodic memory: whereas perceptual details may contribute to the episodic specificity of memory, valence may be tied to more abstract (i.e., less detailed) event representations. Surprisingly, DMN activity was not predicted by the amount of narrative detail recalled, diverging from previous findings implicating lateral temporal and medial frontal regions in narrative retrieval (Baetens et al., 2014; Binder et al., 2009; Gurguryan & Sheldon, 2019). It is possible that the types of narrative details supported by the dorsomedial subnetwork were not captured in participants’ written recalls, as our measure focused on specific story details rather than general conceptual or schematic information. Another possibility is that recall performance was influenced by factors beyond initial memory retrieval, such as forgetting over the one-day delay between scanning and written recall. We found that in-scanner memory ratings strongly related to the number of details recalled the next day, suggesting a meaningful relationship between brain activity and the enduring contents of memory. However, there may have been additional details recalled during the scan that were forgotten during delayed recall, which would not be captured by our analysis approach. Future research incorporating immediate tests and separate measures for general and specific memory content will help clarify how brain activity relates to the persistence of memory details across time.

Past work has identified event-specific patterns of activity in DMN regions, including the medial parietal cortex (Bird et al., 2015) and lateral parietal cortex (Kuhl & Chun, 2014), but less is known about how the maintenance of these patterns is influenced by memory contents. We used two approaches to characterize these patterns: one measuring recall pattern similarity to a high-fidelity video template, and another examining recall pattern stability across repeated retrievals. Emotional valence enhanced recall stability in the dorsomedial subnetwork, while perceptual detail enhanced stability in the medial temporal subnetwork. Comparisons across recalls may capture transformations in the memory representation that distinguish it from perceptual experience (Favila et al., 2020; Xue, 2022). Our recall stability analysis further reveals what event-specific information is maintained consistently across three retrieval rounds. Repeated retrieval can promote a more stable memory trace over time (Antony et al., 2017; Ferreira et al., 2019), accelerating immediate consolidation processes while maintaining the availability of episodic information. Thus, the relationship between recall stability and memory features may reflect early consolidation processes that protect these memory details over time. In contrast, template-recall similarity showed limited association with memory characteristics, suggesting that recall activity patterns may better capture memory quality than the template phase, where other non-remembered perceptual features may influence patterns more strongly.

In conclusion, we identified functional differentiation among DMN subnetworks in supporting memories that persist over time. The dorsomedial subnetwork contributes to an emotional framework for event memory, whereas the medial temporal network is recruited during recall of perceptually detailed memories. By leveraging analyses of both activity and event-specific patterns, the results offer novel evidence for DMN heterogeneity in retrieving the emotional context and perceptual details of past events, informing theoretical accounts of how DMN networks work in tandem to reconstruct complex memory experiences.

## Conflict of interest statement

The authors declare no competing financial interests.

## Acknowledgments

This project was supported by grants from the Brain and Behavior Research Foundation NARSAD Young Investigator Grant (grant number 27663) and National Science Foundation (grant number BCS-2047415). This research was carried out at the Boston University Cognitive Neuroimaging Center, using facilities and equipment supported by the National Science Foundation (grant number BCS-1625552). We would like to thank Stephanie McMains for assistance with MRI data collection, and Zoe Ting, Christina Farmer, Jason Sibrian, and Makayla Romanus for assistance with data scoring.

## Notes

### Competing Interest Statement

The authors have declared no competing interest.

