## Supplemental Materials for "Remembered event features shape default mode network engagement during emotional memory recall"

**Running title:** Default network activity during emotional recall

**Supplemental Materials**

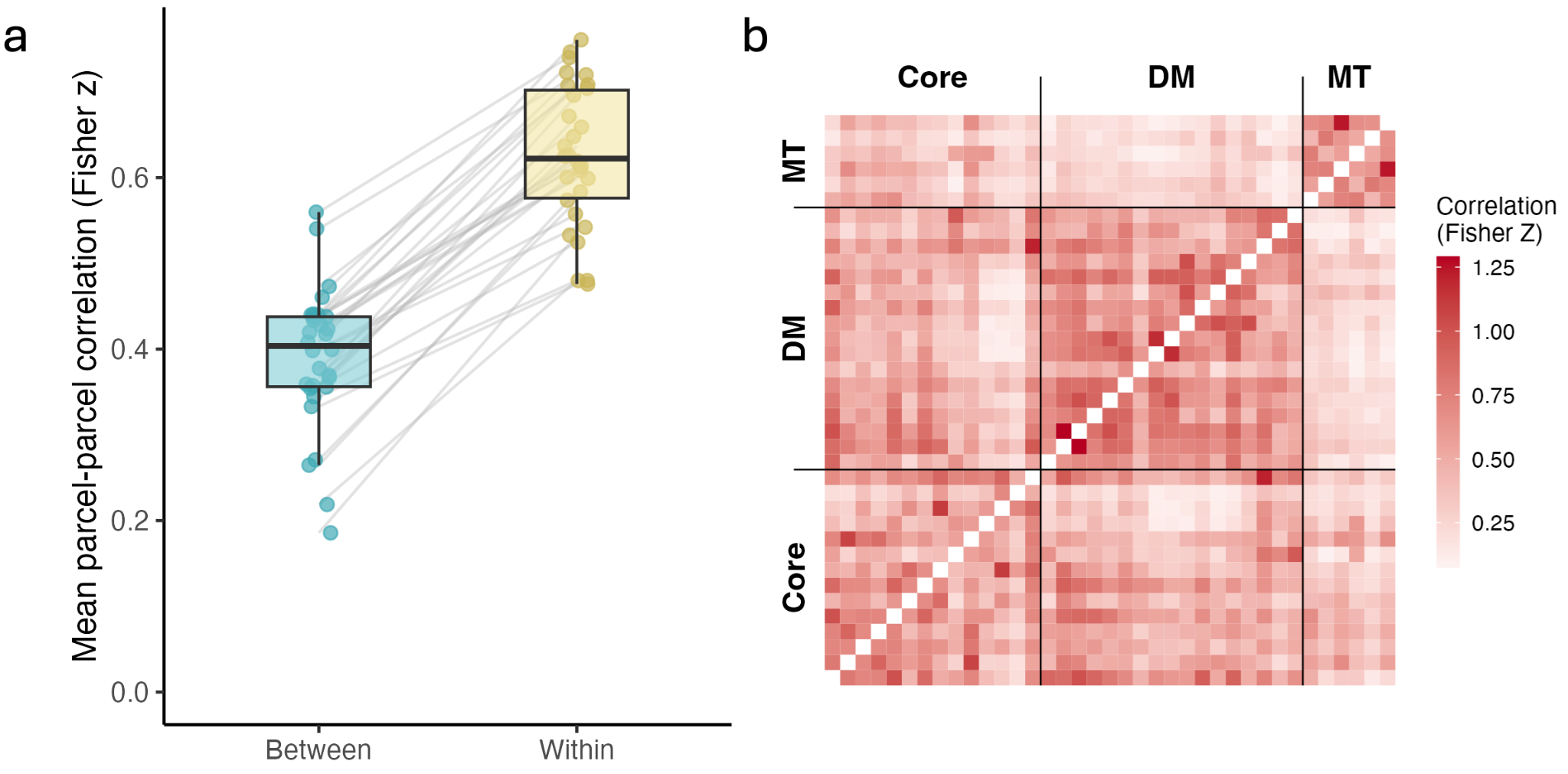

***Supplementary Figure 1.*** **Parcel-level correlations within and between networks.**

**a)** To assess whether network assignments captured meaningful co-variance among parcels in how they were engaged during recall, we computed pairwise correlations of task-related activity (across trials, using the trial averages from our primary analyses). We then tested whether the Fisher z transformed correlations were greater for within-subnetwork parcel pairs than between-subnetwork pairs by averaging within- and between- correlations for each subject and comparing them using a paired-samples t-test. We found that parcel pairs in the same subnetwork showed significantly higher average correlations than those across subnetworks, *t*(29) = 19.735, *p* < .001, indicating strong functional cohesion within each subnetwork during recall.

**b)** Matrix displays the Fisher z-transformed Pearson’s *r* correlation between each parcel defined in the DMN. Black lines indicate separation between the three networks. Darker colors indicate a stronger positive correlation.

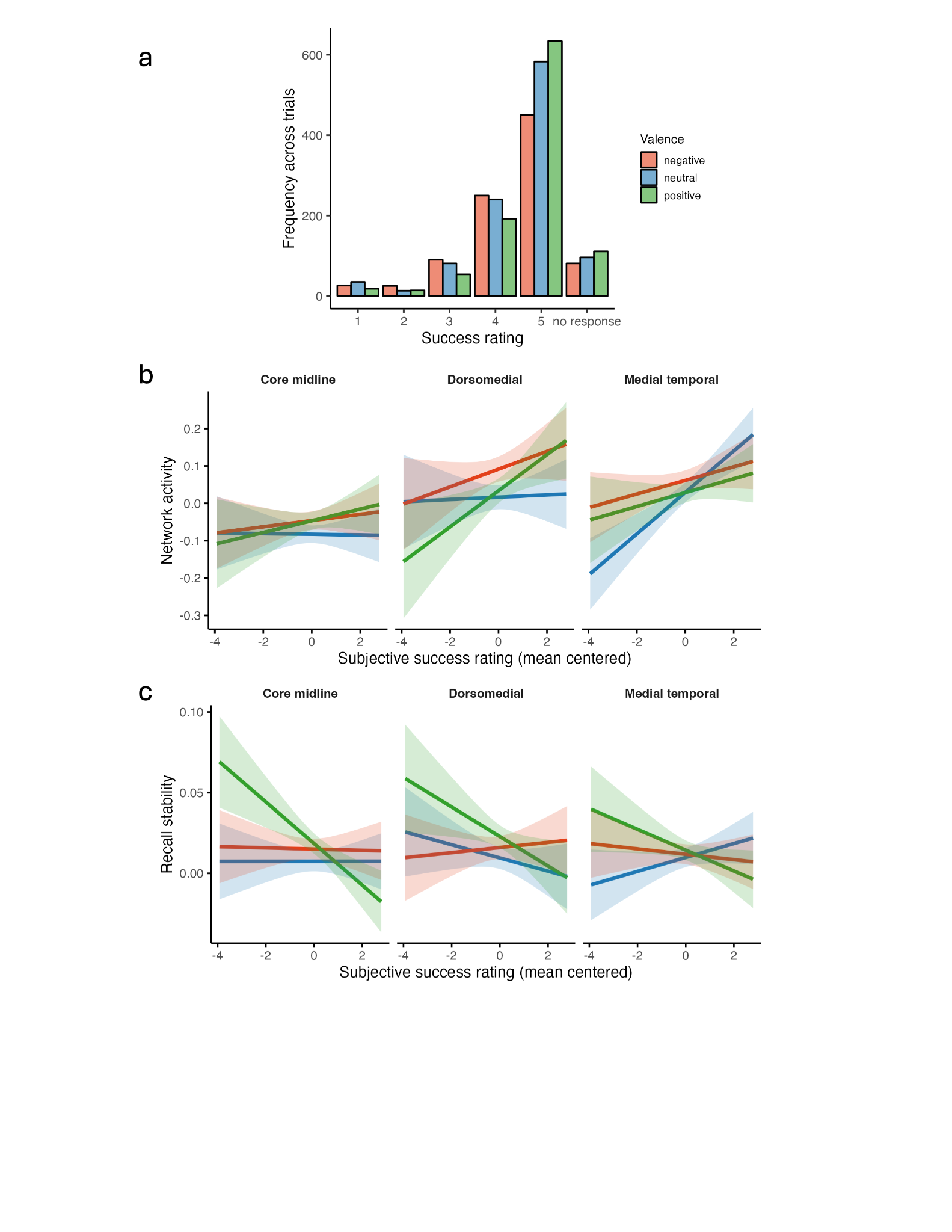

***Supplementary Figure 2. Behavioral characterization and neural correlates of success ratings.***

*a)* Success rating distribution by emotional valence. To characterize participants’ subjective success during encoding, we examined the distribution of in-scanner success ratings. Participants generally reported high levels of success: among trials that were successfully recalled the following day (i.e., those included in the present analyses), 86.8% received a rating of 4 or 5 on the 1-5 success scale. We next tested whether perceived success differed by emotional valence using a linear mixed-effects model predicting success ratings from video valence. Success ratings varied significantly by valence: positive events were rated as more successful than neutral events (*β* = 0.129, *t*(2671.5) = 3.551, *p* < .001), neutral events were rated as more successful than negative events (*β* = -0.129, *t*(2671.8) = -3.486, *p* < .001), and positive events were rated as more successful than negative events (*β* = 0.259, *t*(2672) = 6.983, *p* < .001). Thus, while participants generally reported high success overall, their subjective perception of success varied by emotional valence.

**b)** To investigate how default mode subnetwork activity was related to in-scanner success ratings, we ran a linear mixed-effects model predicting univariate activation from success, valence, and their interaction. The medial temporal subnetwork showed a significant positive association with success ratings (*β* = 0.055, *t*(2473.34) = 4.62, *p_FDR_* < .001), indicating that greater perceived success during in-scanner recall was associated with higher activity in the medial temporal subnetwork. No other DMN subnetwork showed significant effects of success or success-by-valence interactions.

**c)** We conducted an analogous analysis predicting recall stability measures. Across DMN subnetworks, there was no main effect of success ratings on recall stability. However, an interaction was present in core midline and medial temporal subnetwork, such that higher success ratings for positive events were associated with lower recall stability (core midline: *β* = -0.013, *t*(2517.86) = -2.828, *p_FDR_* < .05; medial temporal: *β*= -.011, *t*(2516.69) = -2.540, *p_FDR_* < .05). When the same analysis was conducted using template-recall similarity as the outcome variable, we observed no significant relationships with subjective success.

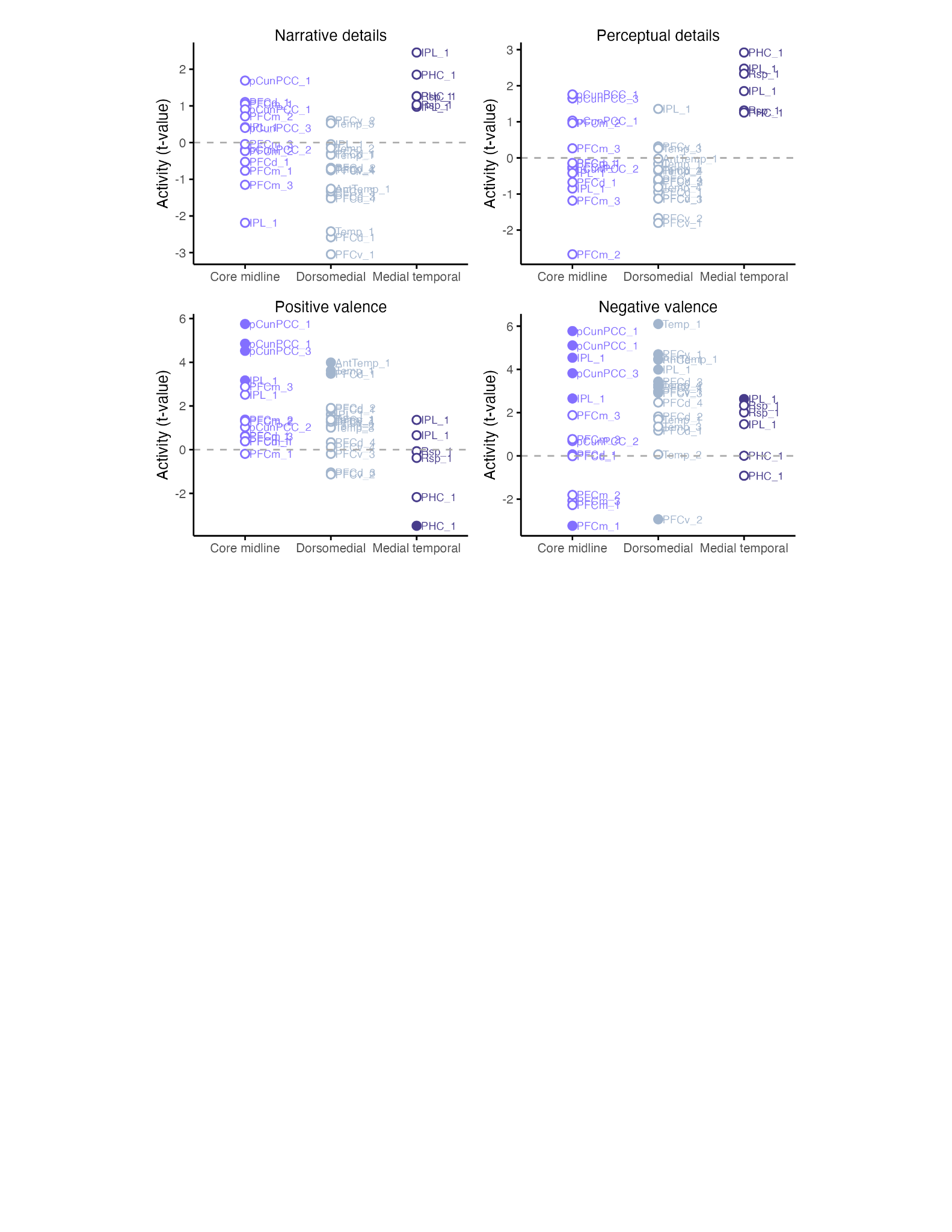

***Supplementary Figure 3.* Activity related to remembered event features: Parcel-level effects.** Plots show t-values from the linear mixed-effects model predicting activity for each individual parcel, with parcels grouped by default mode subnetwork and labeled using Schaefer atlas abbreviations (Schaefer et al., 2018). The corresponding predictor is labeled at the top of each plot. Filled circles indicate parcel-level significance at FDR corrected *p* <.05. We found that valence-related effects were distributed but most prevalent within the core and dorsomedial subnetworks, consistent with our network-level findings. No individual DMN parcel showed significant sensitivity to narrative or perceptual detail. However, visual inspection of parcel-level effects suggests that the medial temporal subnetwork shows coherent recruitment specifically for detail-related effects, suggesting possible subnetwork-level specialization.

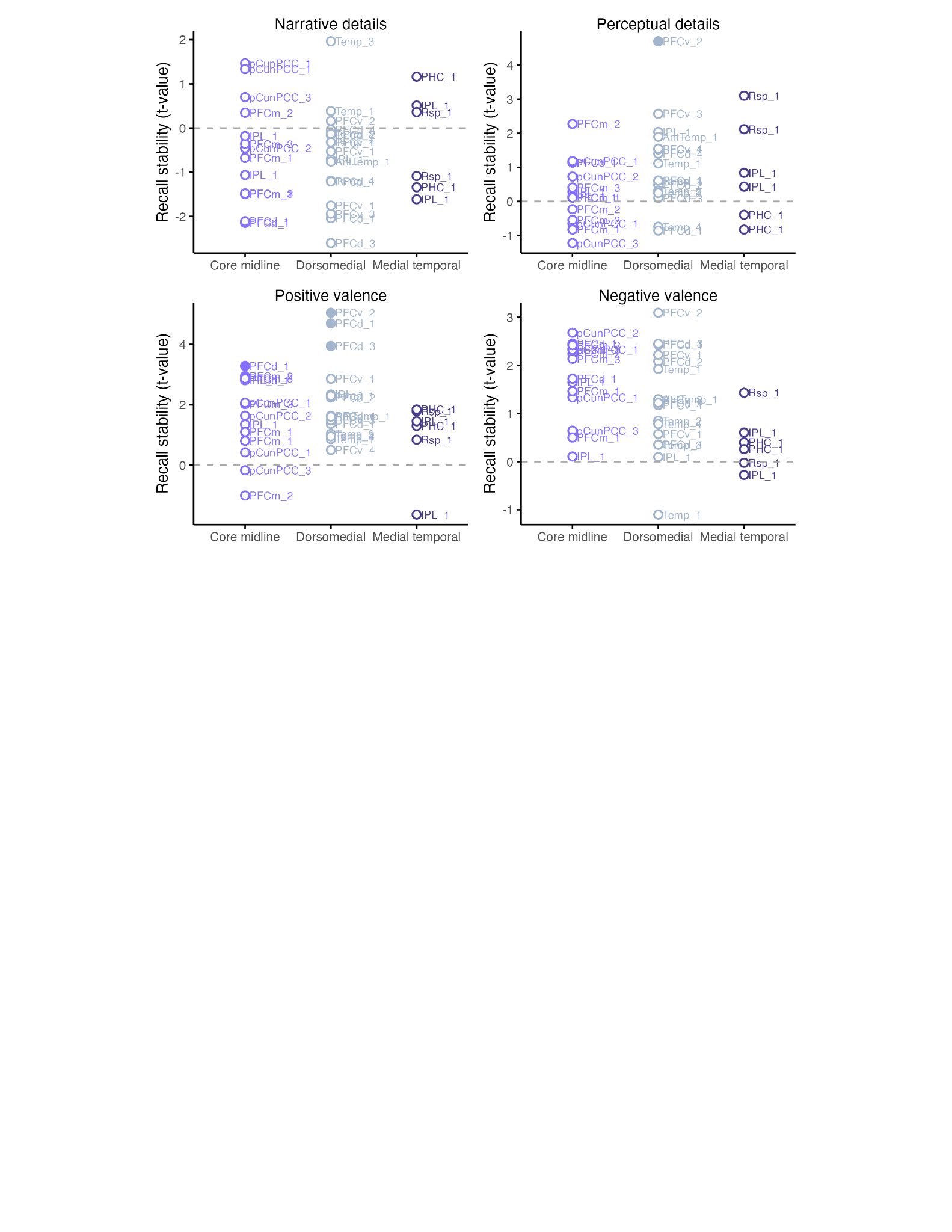

***Supplementary Figure 4.* Recall stability: Parcel-level effects.** Plots show t-values from the linear mixed-effects model predicting recall stability for each individual parcel, with parcels grouped by default mode subnetwork and labeled using Schaefer atlas abbreviations (Schaefer et al., 2018). The corresponding predictor is labeled at the top of each plot. Filled circles indicate significance at FDR corrected *p* <.05. We found that parcels in the dorsomedial and core networks showed greater recall stability for positive memories compared to neutral memories, with a similar trend for negative compared to neutral memories. One region in the dorsomedial network, the ventromedial prefrontal cortex, was also sensitive to perceptual details recalled.

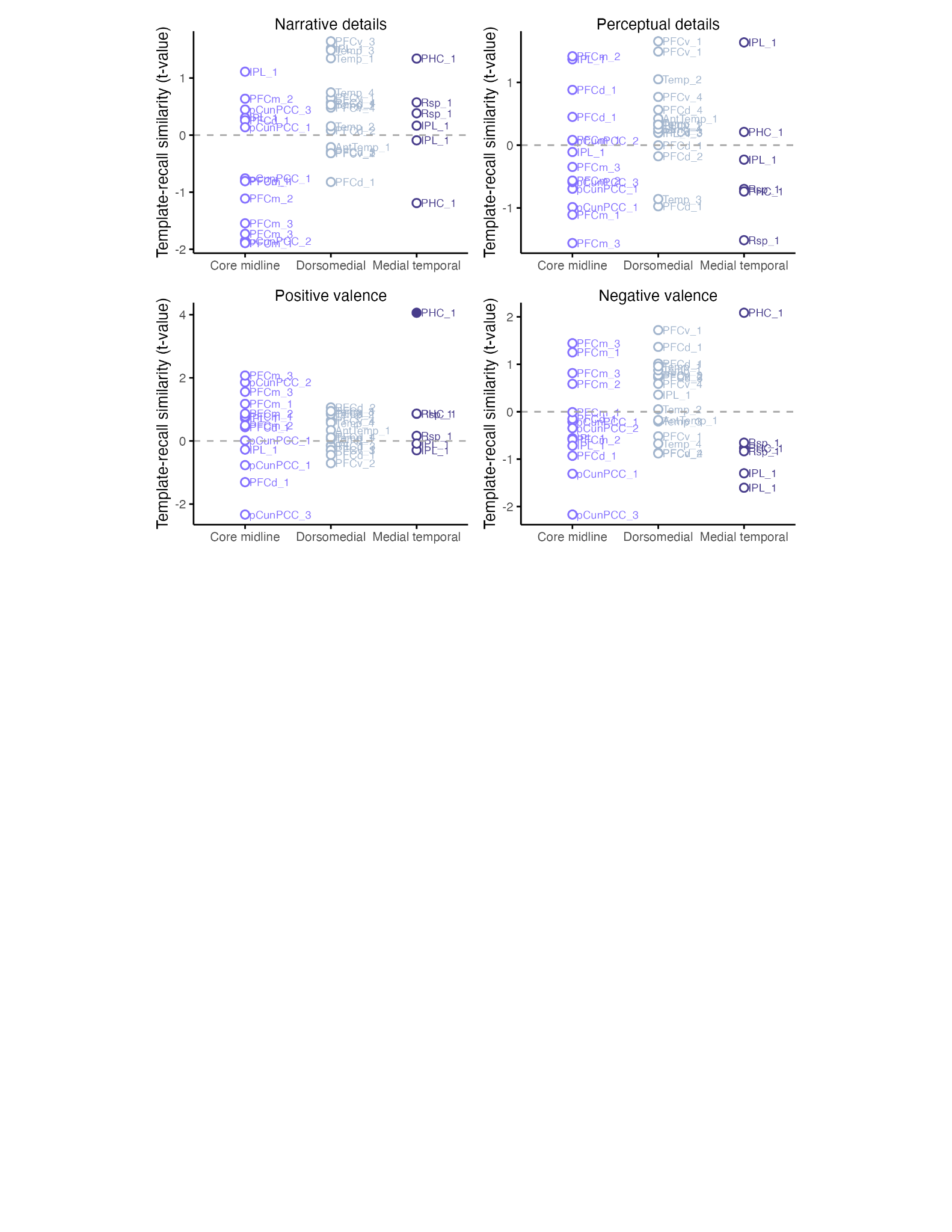

***Supplementary Figure 5.*** **Template-recall similarity: Parcel-level effects.** Plots show t-values from the linear mixed-effects model predicting template-recall similarity for each individual parcel, with parcels grouped by default mode subnetwork and labeled using Schaefer atlas abbreviations (Schaefer et al., 2018). The corresponding predictor is labeled at the top of each plot. Filled circles indicate significance at FDR corrected *p* <.05. Only one parcel showed evidence of being modulated by the features of the remembered videos: the right parahippocampal cortex showed reactivation more for positive than neutral videos, with a similar non-significant pattern for negative videos.

***Supplementary Table 1.*** Significant associations between parcel activity and predictors across the whole brain, using a threshold of FDR-corrected p < .05. Here, DefaultA corresponds to core midline, DefaultB to dorsomedial subnetwork, and DefaultC to medial temporal subnetwork. Network and region labels are from the Schaefer atlas. LH=left hemisphere, RH=right hemisphere.

| **Term** | **Network** | **Region label** | **Hemi** | **Estimate** | **df** | **t** | **p (FDR)** |
| --- | --- | --- | --- | --- | --- | --- | --- |
| Negative valence | ContA | IPS_1 | LH | 0.1380 | 2,720 | 3.25 | p < .01 |
|  | ContA | IPS_1 | RH | 0.1480 | 2,720 | 3.75 | p < .01 |
|  | ContA | PFCl_1 | LH | 0.1200 | 2,720 | 4.62 | p < .001 |
|  | ContA | PFCl_2 | RH | 0.1360 | 2,730 | 6.37 | p < .001 |
|  | ContA | PFCl_3 | LH | 0.0661 | 2,720 | 2.75 | p < .05 |
|  | ContB | IPL_1 | RH | 0.2450 | 2,720 | 4.66 | p < .001 |
|  | ContB | PFCl_1 | LH | 0.1140 | 2,720 | 3.37 | p < .01 |
|  | ContB | PFCld_2 | RH | 0.1160 | 2,720 | 3.92 | p < .01 |
|  | ContB | PFClv_2 | RH | 0.0875 | 2,720 | 2.63 | p < .05 |
|  | ContB | PFCmp_1 | RH | 0.1030 | 2,720 | 4.06 | p < .001 |
|  | ContB | Temp_1 | RH | 0.0562 | 2,720 | 3.84 | p < .01 |
|  | ContC | pCun_1 | LH | 0.1100 | 2,720 | 4.07 | p < .001 |
|  | ContC | pCun_1 | RH | 0.1200 | 2,720 | 4.35 | p < .001 |
|  | ContC | pCun_2 | RH | 0.1240 | 2,720 | 4.94 | p < .001 |
|  | DefaultA | IPL_1 | LH | 0.1040 | 2,720 | 2.65 | p < .05 |
|  | DefaultA | IPL_1 | RH | 0.1160 | 2,720 | 4.54 | p < .001 |
|  | DefaultA | PFCm_1 | LH | -0.0394 | 2,720 | -3.23 | p < .01 |
|  | DefaultA | pCunPCC_1 | LH | 0.0735 | 2,720 | 5.11 | p < .001 |
|  | DefaultA | pCunPCC_1 | RH | 0.0878 | 2,720 | 5.77 | p < .001 |
|  | DefaultA | pCunPCC_3 | LH | 0.0573 | 2,720 | 3.82 | p < .01 |
|  | DefaultB | AntTemp_1 | RH | 0.0608 | 2,720 | 4.48 | p < .001 |
|  | DefaultB | IPL_1 | LH | 0.1100 | 2,720 | 3.99 | p < .001 |
|  | DefaultB | PFCd_1 | RH | 0.1280 | 2,720 | 4.45 | p < .001 |
|  | DefaultB | PFCd_3 | LH | 0.0714 | 2,720 | 3.44 | p < .01 |
|  | DefaultB | PFCv_1 | LH | 0.0499 | 2,720 | 2.99 | p < .05 |
|  | DefaultB | PFCv_1 | RH | 0.1100 | 2,720 | 4.70 | p < .001 |
|  | DefaultB | PFCv_2 | LH | -0.0504 | 2,720 | -2.93 | p < .05 |
|  | DefaultB | PFCv_3 | LH | 0.0978 | 2,720 | 2.91 | p < .05 |
|  | DefaultB | PFCv_4 | LH | 0.0773 | 2,720 | 3.17 | p < .05 |
|  | DefaultB | Temp_1 | RH | 0.1600 | 2,720 | 6.10 | p < .001 |
|  | DefaultB | Temp_4 | LH | 0.0871 | 2,720 | 3.29 | p < .01 |
|  | DefaultC | IPL_1 | RH | 0.0677 | 2,720 | 2.63 | p < .05 |
|  | DorsAttnA | SPL_2 | RH | 0.1310 | 2,720 | 2.74 | p < .05 |
|  | LimbicA | TempPole_2 | RH | 0.0310 | 2,730 | 3.19 | p < .05 |
|  | LimbicB | OFC_1 | RH | 0.0275 | 2,740 | 2.65 | p < .05 |
|  | SalVentAttnA | FrOper_1 | LH | -0.0304 | 2,720 | -2.71 | p < .05 |
|  | SalVentAttnA | ParOper_1 | LH | -0.0921 | 2,720 | -3.32 | p < .01 |
|  | SalVentAttnA | ParOper_1 | RH | -0.0769 | 2,730 | -2.58 | p < .05 |
|  | TempPar | 2 | LH | 0.0638 | 2,720 | 2.62 | p < .05 |
|  | VisCent | ExStr_1 | LH | 0.0528 | 2,720 | 2.68 | p < .05 |
|  | VisCent | ExStr_2 | LH | 0.2000 | 2,720 | 4.46 | p < .001 |
|  | VisCent | ExStr_3 | RH | 0.1420 | 2,720 | 3.46 | p < .01 |
|  | VisCent | Striate_1 | RH | 0.0939 | 2,720 | 2.60 | p < .05 |
| Positive valence | ContB | Temp_1 | RH | 0.0643 | 2,720 | 4.48 | p < .001 |
|  | ContC | Cingp_1 | RH | 0.0397 | 2,720 | 2.98 | p < .05 |
|  | ContC | pCun_2 | RH | 0.1210 | 2,720 | 4.94 | p < .001 |
|  | DefaultA | IPL_1 | RH | 0.0795 | 2,720 | 3.17 | p < .05 |
|  | DefaultA | pCunPCC_1 | LH | 0.0684 | 2,720 | 4.85 | p < .001 |
|  | DefaultA | pCunPCC_1 | RH | 0.0858 | 2,720 | 5.75 | p < .001 |
|  | DefaultA | pCunPCC_3 | LH | 0.0664 | 2,720 | 4.52 | p < .001 |
|  | DefaultB | AntTemp_1 | RH | 0.0531 | 2,720 | 3.99 | p < .01 |
|  | DefaultB | PFCd_1 | RH | 0.0983 | 2,720 | 3.48 | p < .05 |
|  | DefaultB | Temp_1 | RH | 0.0931 | 2,720 | 3.62 | p < .01 |
|  | DefaultC | PHC_1 | LH | -0.0245 | 2,720 | -3.48 | p < .05 |
|  | SalVentAttnA | Ins_1 | RH | 0.0313 | 2,730 | 3.07 | p < .05 |
|  | TempPar | 2 | LH | 0.1070 | 2,720 | 4.48 | p < .001 |

***Supplementary Table 2***. Significant associations between our two RSA measures and predictors across the whole brain, using a threshold of FDR-corrected *p* < .05. Here, DefaultA corresponds to core midline, DefaultB to dorsomedial subnetwork, and DefaultC to medial temporal subnetwork. Network and region labels are from the Schaefer atlas. LH=left hemisphere, RH=right hemisphere.

**Template-recall reactivation**

| **Term** | **Network** | **Region label** | **Hemi** | **Estimate** | **df** | **t** | **p (FDR)** |
| --- | --- | --- | --- | --- | --- | --- | --- |
| Positive valence | DefaultC | PHC_1 | RH | 0.01140 | 2,780 | 4.06 | p < .05 |
| Narrative details × Negative valence | VisCent | ExStr_3 | LH | 0.01560 | 2,790 | 3.84 | p < .05 |
| Perceptual details × Positive valence | VisCent | ExStr_5 | RH | 0.01180 | 2,790 | 3.44 | p < .05 |
|  | VisPeri | ExStrInf_2 | LH | 0.01150 | 2,790 | 4.82 | p < .001 |
|  | VisPeri | ExStrSup_1 | RH | 0.00963 | 2,790 | 3.36 | p < .05 |
|  | VisPeri | StriCal_1 | RH | 0.01350 | 2,790 | 4.57 | p < .001 |

**Recall stability**

| **Term** | **Network** | **Region label** | **Hemi** | **Estimate** | **df** | **t** | **p (FDR)** |
| --- | --- | --- | --- | --- | --- | --- | --- |
| Perceptual details | DefaultB | PFCv_2 | LH | 0.00658 | 2,770 | 4.70 | p < .001 |
|  | DorsAttnB | PostC_4 | RH | 0.00920 | 2,780 | 3.39 | p < .05 |
|  | SomMotA | 9 | RH | 0.00931 | 2,790 | 4.07 | p < .01 |
| Positive valence | ContA | IPS_2 | LH | 0.02920 | 2,770 | 3.64 | p < .05 |
|  | ContB | IPL_1 | RH | 0.03820 | 2,770 | 3.53 | p < .05 |
|  | ContB | PFCld_1 | RH | 0.02370 | 2,770 | 3.07 | p < .05 |
|  | ContB | PFCld_2 | RH | 0.02950 | 2,770 | 3.61 | p < .05 |
|  | ContB | PFClv_2 | RH | 0.02650 | 2,770 | 3.20 | p < .05 |
|  | DefaultA | PFCd_1 | RH | 0.01610 | 2,770 | 3.29 | p < .05 |
|  | DefaultA | PFCm_2 | LH | 0.01930 | 2,770 | 2.95 | p < .05 |
|  | DefaultB | PFCd_1 | RH | 0.03370 | 2,770 | 4.70 | p < .001 |
|  | DefaultB | PFCd_3 | LH | 0.02210 | 2,770 | 3.95 | p < .01 |
|  | DefaultB | PFCv_2 | LH | 0.02230 | 2,770 | 5.05 | p < .001 |
|  | LimbicB | OFC_1 | RH | 0.01090 | 2,770 | 3.54 | p < .05 |
|  | LimbicB | OFC_2 | RH | 0.01040 | 2,770 | 3.22 | p < .05 |
|  | LimbicB | OFC_4 | RH | 0.01820 | 2,770 | 3.01 | p < .05 |
|  | VisCent | ExStr_3 | RH | 0.02320 | 2,780 | 2.96 | p < .05 |
| Narrative details × Negative valence | VisCent | ExStr_3 | LH | 0.01750 | 2,750 | 3.85 | p < .05 |
| Perceptual details × Negative valence | VisPeri | ExStrSup_3 | RH | 0.02170 | 2,790 | 4.37 | p < .01 |
| Perceptual details × Positive valence | ContC | Cingp_1 | RH | -0.00596 | 2,790 | -3.55 | p < .05 |
|  | DorsAttnA | SPL_1 | RH | 0.01150 | 2,790 | 3.52 | p < .05 |
